# Exploring differential interactional preferences of enzyme-bearing dockerins for cohesin domains in the *Clostridium thermocellum* cellulosome

**DOI:** 10.64898/2026.01.19.700288

**Authors:** Snehal Waghmare, Vikas Yadav, Saji Menon, Purnananda Guptasarma

## Abstract

Using its 9 sequentially-placed cohesin (Coh) domains as bait, each CipA scaffoldin chain in a *Clostridium thermocellum* cellulosome binds to (and displays) some combination of 9 out of over ∼70 available dockerin (Doc) domain-bearing lignocellulose-degrading enzymes, at any time. Domains numbered Coh3 through Coh8 show over 94 % pairwise sequence identity with each other, but only ∼ 61-76 % identity with Coh1, Coh2, and Coh9. Such identities are much lower (∼ 40-60 %) amongst Doc domains; however, amongst both Coh and Doc domains, polypeptide backbone folds are highly conserved, suggesting that loading preferences of enzyme-bearing Doc domains upon Coh domains must depend upon their relative abundances, and pairwise affinities. To explore this further, we used microscale thermophoresis (MST), size exclusion chromatography (SEC), native polyacrylamide gel electrophoresis (NPGE), mass spectrometry (MS) and bioinformatics-based approaches (BIBA), to examine 28 Coh-Doc pairwise interactions involving recombinant Coh [Coh1, Coh2, Coh3, Coh9] and enzyme-bearing Doc [Cel8A, Cel9F, Man26/5H, Cel9R, Xyn10C, Xyn11D, Xyn10Z] domains. Interactions were found to occur with varying affinities, suggesting that Coh1 prefers Xyn11D; Coh2 prefers Xyn10C; Coh3 prefers Cel9R; Coh9 prefers Cel9R; Coh1/Coh2 prefer Xyn partners; Coh3/Coh9 prefer Cel partners. Dual modes of binding are shown by Coh1 with Xyn10C and Xyn11D; Coh2 with Xyn10Z, Cel8A, and Cel9R; Coh3 with Cel8A, and Cel9R; and Coh9 with Xyn10C, suggesting that Doc domains use either of their two homologous helices (1 and 3) to bind to Coh domains, as earlier proposed.

**Importance:** Bacteria such as *Clostridium thermocellum* use extracellular enzyme complexes called ‘cellulosomes’ to degrade and use cellulose. Each complex uses a linear chain of nine cohesin (Coh) domains called a ‘scaffoldin’ to bind to (and display) any nine of over seventy available xylan or cellulose-degrading enzymes that bear dockerin (Doc) domains. Understanding interactions between Coh and Doc domains facilitates an appreciation of how cellulosomes are assembled and supports the building of protein-engineered constructs that utilize such interactions for many conceivable enzymatic and other applications. The significance of the presented research lies in its demonstration of the differential modes and pairwise affinities of different Coh-Doc interactions, using recombinant protein constructs and a combination of quantitative, semi-quantitative and qualitative analytical methods.

## Introduction

The term “cellulosome” was first used to denote an extracellular, high molecular weight, multi-enzyme complex produced by the anaerobic bacterium, *Clostridium thermocellum*, a microbe well known for its ability to degrade and use the saccharide components of cellulose as a source of nutrition; in particular, carbon [1]. The cellulosome functions like a multi-modular, multi-functional, nanoscale machine that degrades plant cell wall polysaccharides such as xylan and cellulose into small saccharides that are then imported and used by *C. thermocellum* [2–3]. At its core, the cellulosome hosts a protein called a scaffoldin, which functions as a receptable for enzymes, and consists of a multi-domain polypeptide chain called CipA which contains nine contiguously-placed type I ‘cohesin (Coh)’ domains, numbered Coh1 to Coh9 [1–3]. Surrounding CipA, and bound to the nine Coh domains of CipA, is an assortment of enzymes containing type I ‘dockerin (Doc)’ domains. Enzymes use these Doc domains to recognize and bind to CipA’s Coh domains. Each Doc domain is a sequence variant of every other Doc domain [4]. Currently, seventy-two different enzymes encoded by the *C. thermocellum* genome, such as cellulases (endoglucanases, exoglucanases, and glucosidases), or hemicellulases (xylanases, or lichenases) are known to possess a Doc domain that could conceivably bind to a a CipA chain through one of its Coh domains [5–12]. This suggests that (i) any nine of the seventy-two available enzymes could be immobilized by each CipA chain at any point of time, and also that (ii) different CipA chains could host different sets of nine enzymes, depending upon how many CipA chains are available in respect of the available Doc domain-bearing enzymes, and their relative availabilities. In addition to its nine Coh domains, each CipA chain also contains (a) a carbohydrate binding module (CBM) domain, positioned between domains Coh2 and Coh3, as well as (b) a type II Doc domain located at the C-terminus of the CipA chain, to facilitate the binding of CipA to bacterial cell surface proteins such as SdbA, Orf2p, OlpB, and OlpA, which host a type II Coh domain capable of binding to a type II Doc domain [13].

Therefore, it is clear that *C. thermocellum* displays a multitude of CipA chains, and that each of these very likely displays nine of the bacterium’s seventy-two different cellulose-degrading enzymes. This description evokes the image of a cellulosome being like a multi-capability ‘swiss knife’ since each enzyme has a slightly different glycosidic bond-hydrolysing function, with the difference being that different enzymes bound to any CipA chain are like birds sitting on a wire which can dissociate and reassociate to alter the sequence, and composition, of enzyme display by the CipA chain. In turn, this suggests that enzyme-bearing Doc domains must become loaded upon Coh domains in a manner that is dependent upon both (A) the relative abundances of enzyme-bearing Doc domains and Coh domain-bearing CipA chains, and (B) the pairwise affinities amongst the different Coh and Doc domains. However, such pairwise affinities have not been studied in detail to establish the extent to which they differ. This is the lacuna that is addressed by this paper. We address both the larger question of whether indeed any enzyme-borne Doc domain can bind to any CipA-borne Coh domain, in principle, and also the smaller and more detailed question of whether indeed binding preferences could determine which enzyme would locate upon which Coh domain on the CipA chain.

Doc domains are ∼60-70 residues in length, and folded into all α-helical structures, with chains containing two duplicated copies of a 22 residues-long polypeptide stretch [14–16]. Doc domains serve no catalytic function, and exist in many different enzymes [17–18]. Calcium plays an important role in Doc domain stability and function, and it has been demonstrated that the chelation-based removal of calcium by EDTA renders Doc domains incapable of interacting with Coh domains [16, 19]. Coh domains, in contrast, are longer and range in length from ∼140-150 residues, and are folded into beta sheets, using a nine-stranded β-barrel with an overall “jelly roll” topology stabilized by a conserved hydrophobic core [20]. The affinities of Type I Coh domains for Type I Doc domains are believed to be amongst the strongest of affinities known to protein biochemistry, with dissociation constants approaching 10^-9^ M for some instances that have been studied [21–23], and with evidence pointing towards such high affinities translating into extended periods of association amongst Coh and Doc domains [2, 20, 24]. The binding interface between Coh and Doc domains is primarily characterized by hydrophobic interactions, supplemented by hydrogen-bonding [25]. Structural studies of Coh-Doc complexes indicate that Coh and Doc domains undergo no significant conformational change upon binding to each other [26], suggesting that Coh-Doc interactions are cases of ‘rigid body docking’.

In the state of interaction with a Coh domain, a Doc domain typically binds through one of its two α-helix repeats, i.e., using helix 1 or helix 3 [25]. Thus, a generalized ‘dual-binding mode’ for Type I Coh-Doc interactions has been proposed in which helix 1, or 3, bind to the Coh domain, through a relative rotational reorientation of 180° of the Doc domain in respect of the Coh domain [25, 27]. This suggests that, in a large population of interacting species, any Type I Doc domain can potentially bind to any Type I Coh domain in either or both orientations, but also that for singular instances of binding, this might occur in a mutually-exclusive fashion, since the high affinity of interaction would prevent dissociation and reorientation events. Both helix 1 and helix 3 of a Doc domain contain a conserved pair of adjacent serine (Ser) and threonine (Thr) residues which establish a hydrogen-bonding network that facilitates binding [25, 28]. Our opinion is that the existence of two binding modes provides scope for improved quaternary structural interactions inside densely-packed multienzyme complexes: Whenever a Doc domain is unable to bind to an otherwise-unoccupied Coh domain upon a CipA chain in one mode, for a purely steric reason, i.e., because the attached enzyme gets in the way of something else that is already bound, the enzyme can instead bind to the same unoccupied Coh domain using the other binding mode. The alternative mode of binding places the attached enzyme on the other side of the linear CipA chain, due to the 180° rotational reorientation. In this manner, alternative binding modes probably facilitate the approaches of different enzymes of different sizes towards a CipA chain, in respect of other vicinal and flanking enzymes that are pre-bound to the same CipA chain. It must also be noted that although the above-mentioned Ser-Thr motifs exist in all Doc domains of *C. thermocellum*, there are notable variations in their sequence residue contexts, and variations in neighbouring residues can potentially influence their interactions with Coh domains. Such variations have indeed been speculated to be the basis for the potential existence of distinct ligand-binding specificities and interactional preferences between Coh and Doc domains, which can determine the relative positioning of different enzymes upon CipA chains [29]. Together with the interactional flexibility afforded by the dual-binding mode, this can facilitate reorganization of enzymes upon cellulosomes, through steric interactions between neighbouring enzymes that jostle for space upon CipA.

Challenges in the production of recombinant proteins consisting of only Doc domains (i.e. without the attached enzyme) have arisen due to low expression levels, poor solubility, and degradation, preventing comprehensive binding analyses [5, 30–31]. Despite this, Coh-Doc interactions have been qualitatively and quantitatively investigated in *C. thermocellulm* and other species of *Clostridium* such as *C. josui*, and *C. cellulolyticum*. These investigations have suggested that subtle differences in tertiary micro-structural surface features (determined by differences in sequence) may be responsible for some amount of specificity in Coh-Doc interactions. Clearly, further analyses of intra-species and inter-species Coh-Doc complexes will reveal molecular determinants of specificity and selectivity in Coh-Doc interactions [29, 32–33]. Intriguingly, the Ser-Thr motif conserved in Type I enzyme-bearing Doc domains of *C. thermocellum* is absent in analogous domains of *C. cellulolyticum*, where hydrophobic residues replace the Ser-Thr sequence motif, indicating how nature creates species-specific interaction schemes [34]. Substitution of the Thr residue in helix 1 of a *C. thermocellum* enzyme-bearing Doc domain by a Leu residue enables its recognition by a Type I *C. cellulolyticum* Coh domain [35]. Studies also indicate that binding specificity, even in *C. thermocellum* Doc domains, may be governed by the entire surfaces of helices 1, or 3, and not just by the Ser-Thr motif [36]. Lytle et al. 1996, studied the interaction of the Doc domain of CelS with four cohesins (Coh 1, 2, 3 and 9). The differences in the Coh domains apparently had no effect on their abilities to bind to the Doc domain of CelS. It is reported that although subtle differences in affinity might exist, thioredoxin-fused Doc domain of CelS bind to Coh domains of CipA non-selectively, to form stable complexes, independent of Coh domain identity [37]. Given the above background, and pending questions about Coh-Doc interactions, we describe below results from quantitative, semi-quantitative, and qualitative analyses of a limited set of *C. thermocellum* Coh-Doc interactions using recombinant Coh domains and recombinant enzyme-bearing Doc domains, to discuss how these results reconcile current conceptions of non-selectivity/non-specificity of intra-species Coh-Doc interactions with intra-species preferences of certain enzyme-bearing Doc domains for certain Coh domains.

## Results

### Cohesin (Coh) and Dockerin (Doc) domains used: Structure-aligned sequence comparisons

We begin with an analysis of the conservation of residues at different positions in the known sequences of various Coh and Doc domains, across cellulosomes from different bacterial species. A few Coh-Doc complexes from *C. thermocellum* have been crystallized. Figure 1 shows the known structure of one such complex (PDB ID: 1OHZ), overlayed with a color-based representation of residue conservation, using Consurf software. From the figure, considerable sequence conservation is evident in the beta sheet-based Coh domains, as well as in the alpha helix-based Doc domains, with especially high sequence conservation in regions directly involved in Coh-Doc interactions at the interface of the two domains.

**Figure 1:**
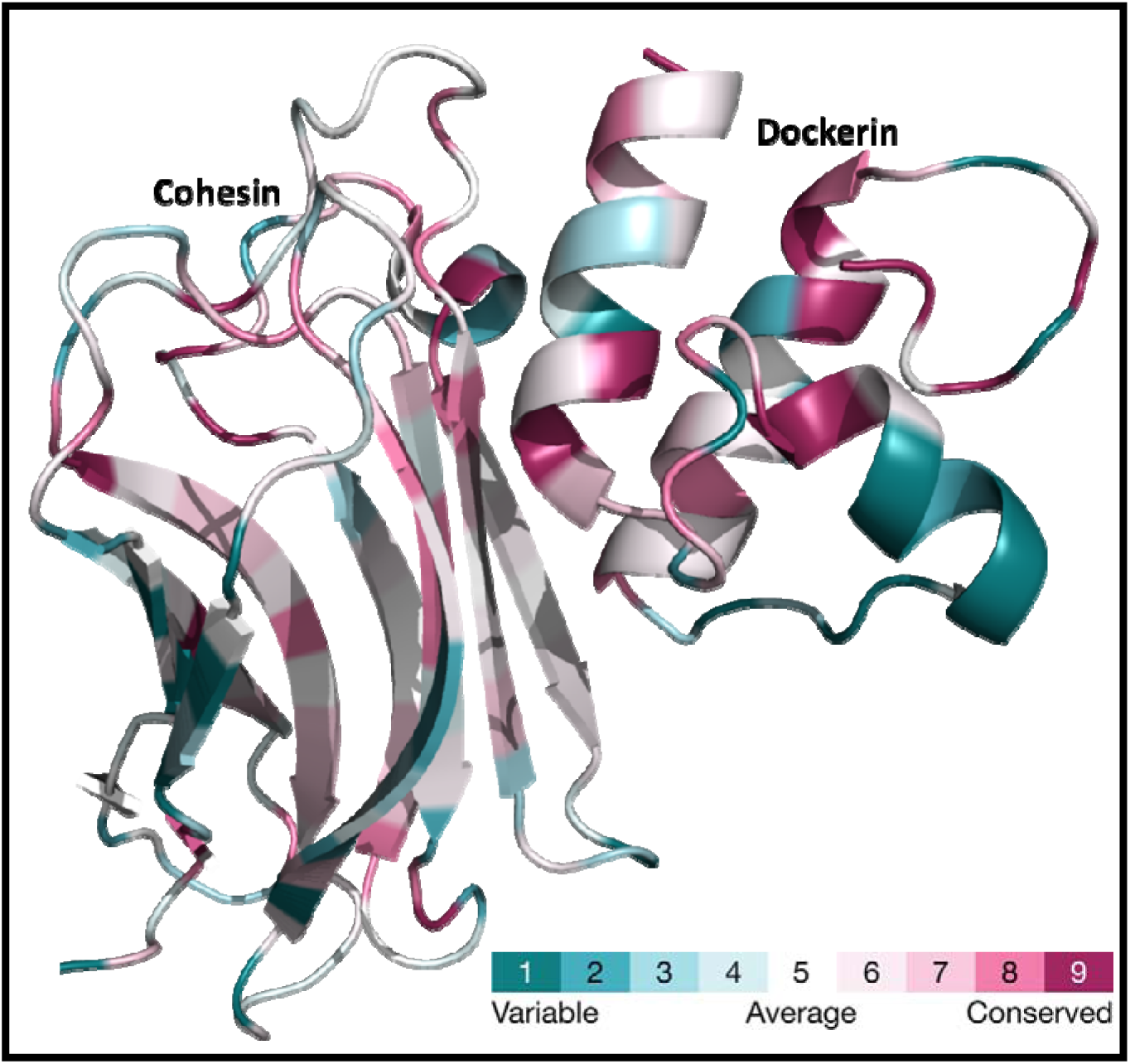
ConSurf analysis showing sequence conservation across bacterial species in different regions of Coh and Doc domains, overlayed on the structure of the Coh-Doc complex in PDB ID: 1OHZ.

Supplementary Figures 1, 2, 3, 4 and 5, respectively, present (1) a sequence alignment of the nine Coh domains of CipA; (2) a matrix that lists the percent identities for all pairwise sequence alignments amongst CipA’s Coh domains; (3) a sequence logo representation of residue occurrence frequency at different sequence positions in the nine Coh domains, generated using Weblogo 3; (4) a similar sequence logo representation for only the four Coh domains selected for the study presented in this paper (Coh1, Coh2, Coh3 and Coh9), eliminating domains that are either identical, or almost identical, to these four domains; and (5) amino acids present in the strands of each beta sheet of the four selected Coh domains, noting that interactions with Doc domains involve only strands 5, 6, 3 and 8. From the said Supplementary Figures, it is evident that the nine Coh domains in the CipA scaffoldin of *C. thermocellum* share more than 90 % sequence identity, and that domains 4 and 5 are 100 % identical to each other, as are domains 6 and 8, indicating the occurrence of a duplication of regions within the CipA gene encoding one Coh domain corresponding to each of these two pairs of identical domains, during evolution.

In a similar vein, Supplementary Figures 6, 7 and 8, respectively, show (6) a sequence alignment of Doc domains present in seven enzymes selected for this study (CelA, CelF, CelH, CelR, XynC, XynD, and XynZ); (7) a matrix that lists the percent identities for all pairwise sequence alignments of these seven Doc domains; and (8) a sequence logo representation of residue occurrence frequency at different sequence positions in these seven Doc domains, generated using Weblogo 3. From Supplementary Figure 7, it is evident that Doc domains show roughly ∼50 % sequence identity with each other. Supplementary Figure 6 shows that helices 1 and 3 are nearly identical, explaining why either of these helices appears to be capable of potentially engaging in interactions with Coh domains, and perhaps even both of these helices, for a particular Doc domain, albeit in a mutually-exclusive manner for a single copy of such a Doc domain. In each of helices 1 and 3, it may be noted that there is a pair of next-neighbour serine (S) and threonine (T) residues that is thought to be directly involved in Coh-Doc interactions.

### Coh domains and enzyme-bearing Doc domains produced: Structure and stability comparisons

This section describes the characteristics of Coh domains and enzyme-bearing Doc domains used for pairwise interaction studies. Figures 2A and 2B, respectively, show the SDS-PAGE profiles of individual Coh domains, and enzyme-bearing Doc domains produced, demonstrating acceptable levels of purification. Figures 2C and 2D, respectively, show circular dichroism (CD) spectra of a representative Coh domain (Coh2), and a representative enzyme-bearing Doc domain (XynD), demonstrating beta sheet content in the Coh domain and a mixed beta-alpha structure in the enzyme-bearing Doc domain. Figures 2E and 2F, respectively, show differential scanning calorimetric (DSC) thermograms demonstrating the unfolding of folded protein entities, and establishing the temperatures of melting (T_m_s) of Coh2, and XynD. Supplementary Figures 9, 10, 11 and 12, respectively, present the CD spectra of all the other Coh domains, the DSC thermograms of all the other Coh domains, the CD spectra of all the other enzyme-bearing Doc domains, and the DSC thermograms of all the other enzyme-bearing Doc domains shown to be produced and purified in Figures 2A and 2B. Further, Supplementary Figure 13 presents tryptophan fluorescence data in support of the enzyme entities in the enzyme-bearing Doc domains displaying cooperative unfolding transitions upon heating. Since the Coh domains possess no tryptophan residues, no fluorescence experiments were performed to monitor unfolding transitions in these domains.

**Figure 2:**
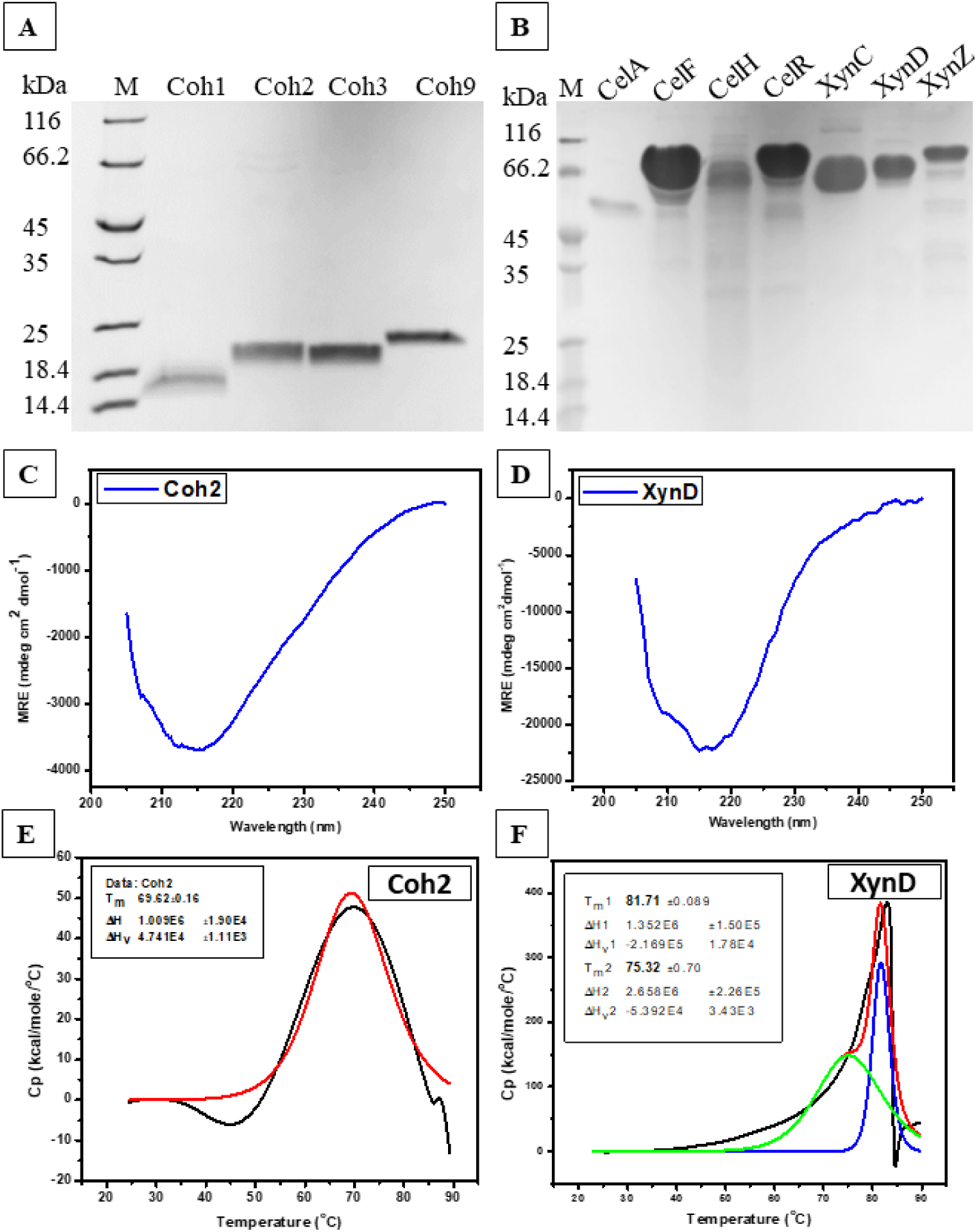
A) SDS-PAGE of all selected Coh domains; Lane 1: Protein marker; Lane 2: Coh1 (17.13 kDa); Lane 3: Coh2 (16.54 kDa); Lane 4: Coh3 (17.32 kDa); Lane 5: Coh9 (17.25 kDa). B) SDS-PAGE of all natural enzyme-bearing Doc domains; Lane 1: Protein marker; Lane 2: CelA (50.2 kDa); Lane 3: CelF (82.1 kDa); Lane 4: CelH (99.04 kDa); Lane 5: CelR (80.9 kDa); Lane 6: XynC (67.63 kDa); Lane 7: XynD (70.2 kDa); Lane 8: XynZ (90.64 kDa). C) Circular dichroism profile of Coh2. D) Circular dichroism profile of XynD. E) DSC thermogram of Coh2. F) DSC thermogram of XynD.

Together, all of these pieces of data establish that each of the entities produced is stably folded and, therefore, competent to be subjected to further investigations of pairwise interactions amongst Coh and enzyme-bearing Doc domains. The data for the Coh domains shows that they are quite stable in their folded states (as independently-produced entities), displaying single and cooperative thermal unfolding transitions with T_m_ values ranging from 61 □ to 72 □. The enzyme-bearing Doc domains typically showed two unfolding transitions in the neighborhood of 75 □ to 88 □, which could owe to the independent unfolding of the catalytic domain, the carbohydrate-binding module (CBM; present in six of the seven enzymes produced), or the Doc domain that exists in fusion with these domains. It is not possible to determine from the unfolding data itself exactly which pair of domains displays unfolding independently of the others, without performing control experiments in which each domain is produced as an independent entity. However, we may note that (a) the data clearly establishes that the enzyme-bearing Doc domains exist as very stably-folded entities, and (b) the likelihood is high that the Doc domain unfolds independently of the enzyme and any CBM domain present, since the independence of movement of the Doc domain (to facilitate assembly upon CipA) could cause it to unfold independently in a DSC experiment.

### Quantitative analyses of cohesin-dockerin interactions by microscale thermophoresis (MST)

We also performed isothermal titration calorimetry (ITC) experiments, but these could not be reliably interpreted, owing to (i) our inability to distinguish heat of binding from heat of dilution for the low protein concentrations we were able to achieve, (ii) the difficulty of distinguishing amongst different modes of binding (data not shown). Therefore, we shifted our efforts to using microscale thermophoresis (MST) experiments. Interactions were allowed to occur between (A) a fixed concentration of an independently-expressed and purified Coh domain, e.g., one out of Coh1, Coh2, Coh3, or Coh9, labelled with the dye red-NHS, and (B) varying concentrations of an enzyme-bearing Doc domain, e.g., one out of the seven independently-expressed enzymes, CelA, CelF, CelH, CelR, XynC, XynD, or XynZ, within a series of solutions that were then transferred into a series of capillaries. Designated regions of all of these capillaries were simultaneously heated and then subsequently cooled, and the rates of depletion and restoration of fluorescence signal due to mass transport of the labeled cohesin (to cooler regions during heating, or back to the original regions during cooling) were monitored, with or without bound enzyme-bearing Doc domain. The plots of fraction of cohesin domain bound are shown in Figure 3, as a function of varying concentrations of the ligand (enzyme-bearing Doc domain, plotted on a semi-logarithmic scale), from which dissociation constants (K_d_ values) were determined. A representative set of plots is shown for interactions between Coh9 and each of seven enzyme-bearing Doc domains. Supplementary Figures 14, 15 and 16, respectively, show similar plots for interactions of all seven enzyme-bearing Doc domains with Coh1, Coh2, and Coh3. In all of these plots, the presence of a single curve indicates a monophasic nature of binding, while the presence of two curves indicates a biphasic nature of binding (potentially corresponding to two modes of binding of enzyme-bearing Doc domains to a Coh domain, using either helix 1, or helix 3, on the Doc domain). Notably, where a single curve was obtained, instead of two, this could indicate either that (i) binding is dominated by one helix (helix 1 or helix 3) or, alternatively, that (ii) binding using both helices occurs with comparable affinities, causing both to contribute to the same binding curve. In contrast, where two curves convertible to two different K_d_ values were obtained, it would appear that one mode of binding was weaker than the other.

**Figure 3:**
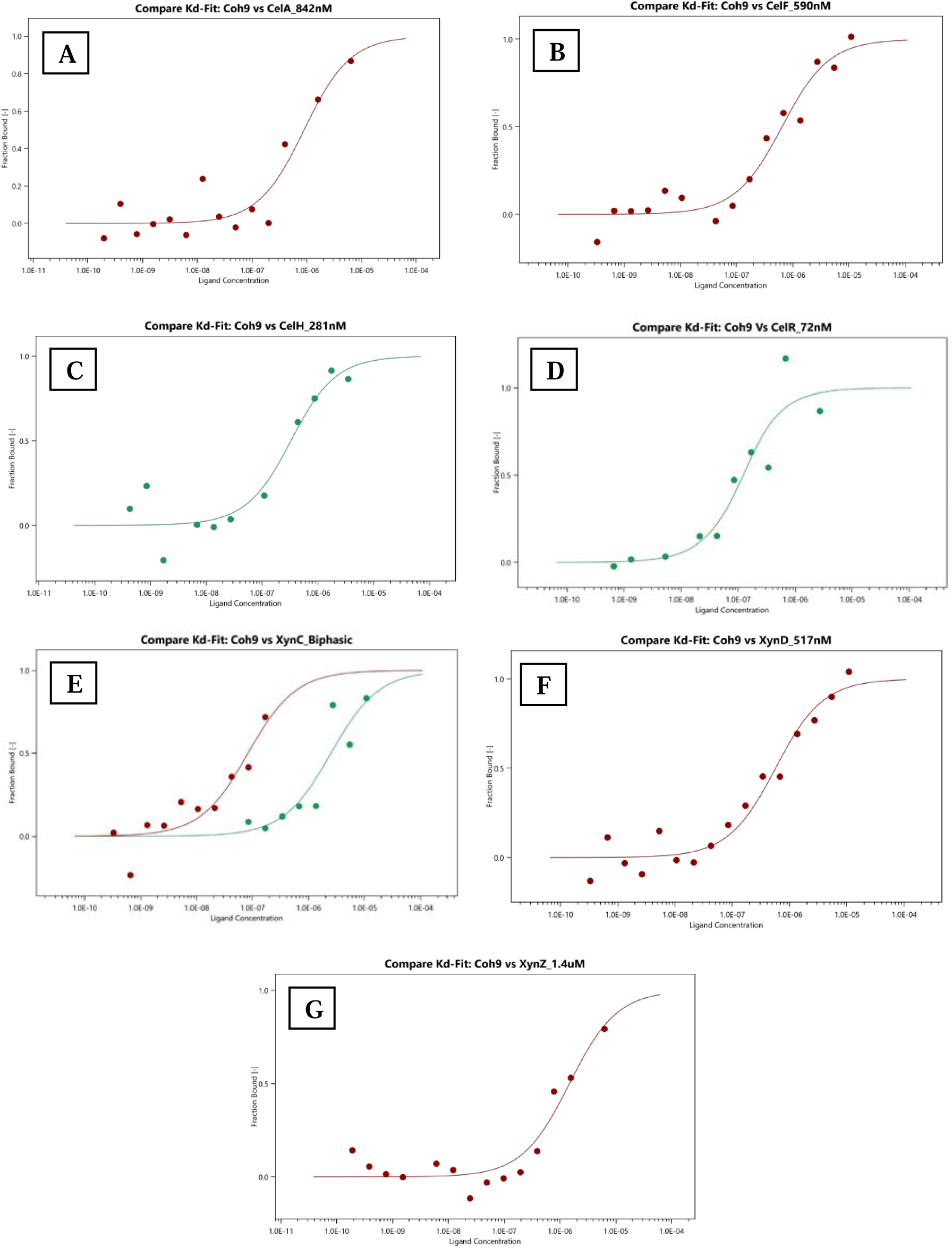
Quantitative interaction analysis of Coh9 with dockerins of selected enzymes using micro scale thermophoresis. (A) CelA (B) CelF (C) CelH (D) CelR (E) XynC (F) XynD and (G) XynZ.

Table 1 summarizes the estimated K_d_ values for all twenty-eight pairwise interactions. Analyses of these suggests that (a) Coh1 binds in a monophasic manner to Doc domains associated with all four cellulases, but in a biphasic manner to Doc domains associated with two out of three xylanases (XynC and XynD); displaying the strongest interaction with the Doc domain on XynC, with a K_d_ of ∼1.0 nM; (b) Coh2 binds in a biphasic manner to the Doc domains associated with two cellulases (CelA and CelR) and one xylanase (XynZ), but in a monophasic manner with the Doc domains associated with all other enzymes; binding most strongly to XynC, with a K_d_ of ∼4.4 nM; (c) Coh3 binds in a biphasic manner to the Doc domains associated with two cellulases (CelA and CelR), and in a monophasic manner to the Doc domains associated with all other enzymes; binding most strongly to CelR, with one mode of binding involving a K_d_ of ∼1.54 nM, and an additional weaker mode of binding involving a K_d_ of ∼760 nM; (d) Coh9 binds in a biphasic manner to the Doc domain associated with a xylanase (XynC), and in a monophasic manner to the Doc domains associated with all other enzymes; binding most strongly to CelR, with a K_d_ of ∼72 nM.

**Table 1:**
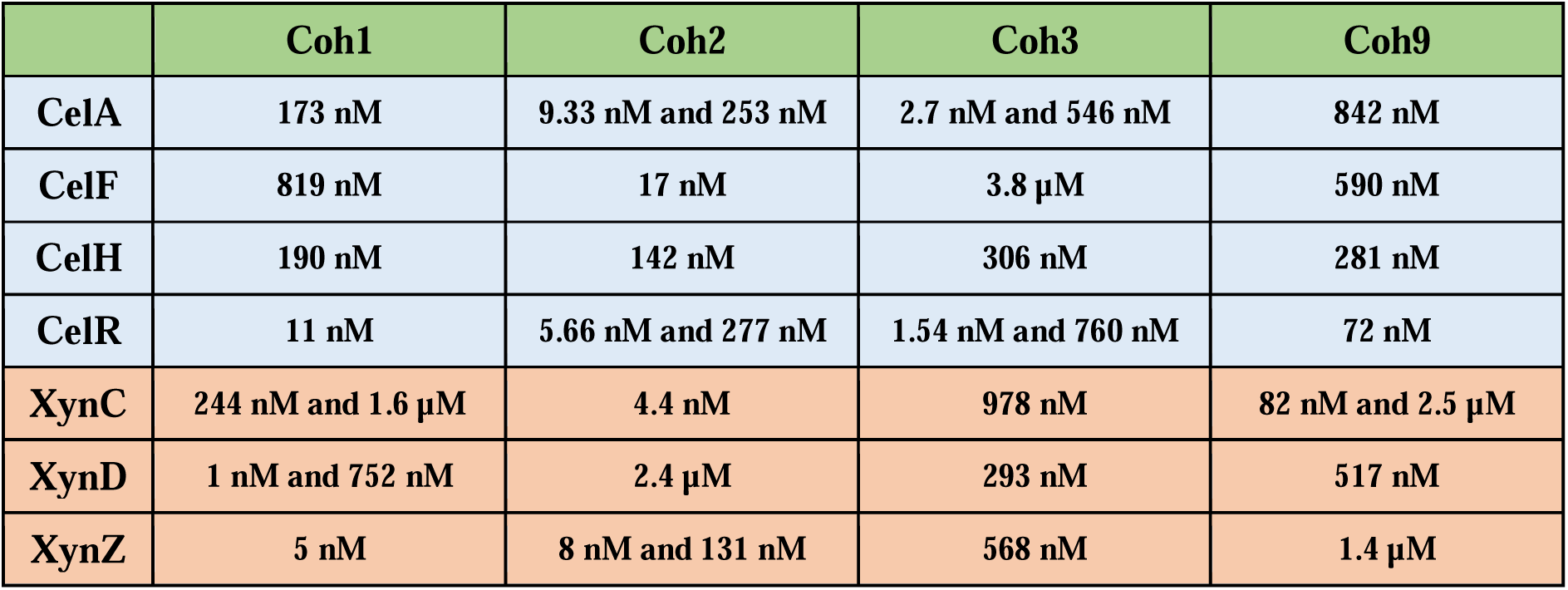
K_d_s estimated for pairwise cohesin-dockerin interactions using microscale thermophoresis.

|  | Coh1 | Coh2 | Coh3 | Coh9 |
| --- | --- | --- | --- | --- |
| **CelA** | 173 nM | 9.33 nM and 253 nM | 2.7 nM and 546 nM | 842 nM |
| **CelF** | 819 nM | 17 nM | 3.8 μM | 590 nM |
| **CelH** | 190 nM | 142 nM | 306 nM | 281 nM |
| **CelR** | 11 nM | 5.66 nM and 277 nM | 1.54 nM and 760 nM | 72 nM |
| **XynC** | 244 nM and 1.6 μM | 4.4 nM | 978 nM | 82 nM and 2.5 μM |
| **XynD** | 1 nM and 752 nM | 2.4 μM | 293 nM | 517 nM |
| **XynZ** | 5 nM | 8 nM and 131 nM | 568 nM | 1.4 μM |

### Semi-quantitative analyses of Coh-Doc interactions by size exclusion chromatography (SEC)

We next used size exclusion chromatography (SEC) to monitor the elution of protein entities from a 24 ml bed-volume SEC column, separating three different populations, namely: (i) the population of an enzyme-bearing Doc domain, eluting in the absence of a bound Coh domain (at ∼14-15 ml); (ii) the population of a Coh domain, eluting in the absence of a bound enzyme-bearing Doc domain (at ∼17 ml); and (iii) the population of complexes formed between a Coh domain and an enzyme-bearing Doc domain (at < ∼14-15 ml). Elutions were monitored through measurement of absorption of ∼280 nm light, bearing in mind that Coh domains absorb ∼280 nm light poorly (due to their lack of tryptophan and content of only 3 to 5 tyrosine residues which absorb relatively poorly at ∼280 nm). Areas covered by peaks corresponding to free enzyme-bearing Doc domains, free Coh domains, and complexes of Coh and enzyme-bearing Doc domains, were used to infer trends of interactions from relative areas of peaks corresponding to unbound enzymes and Coh-bound enzyme-bearing Doc domains. Figures 4 and 5 show SEC profiles for experiments involving different Coh domains and all seven enzyme-bearing Doc domains. The overall preferences inferred from elution area analyses are summarized in Table 2.

**Figure 4:**
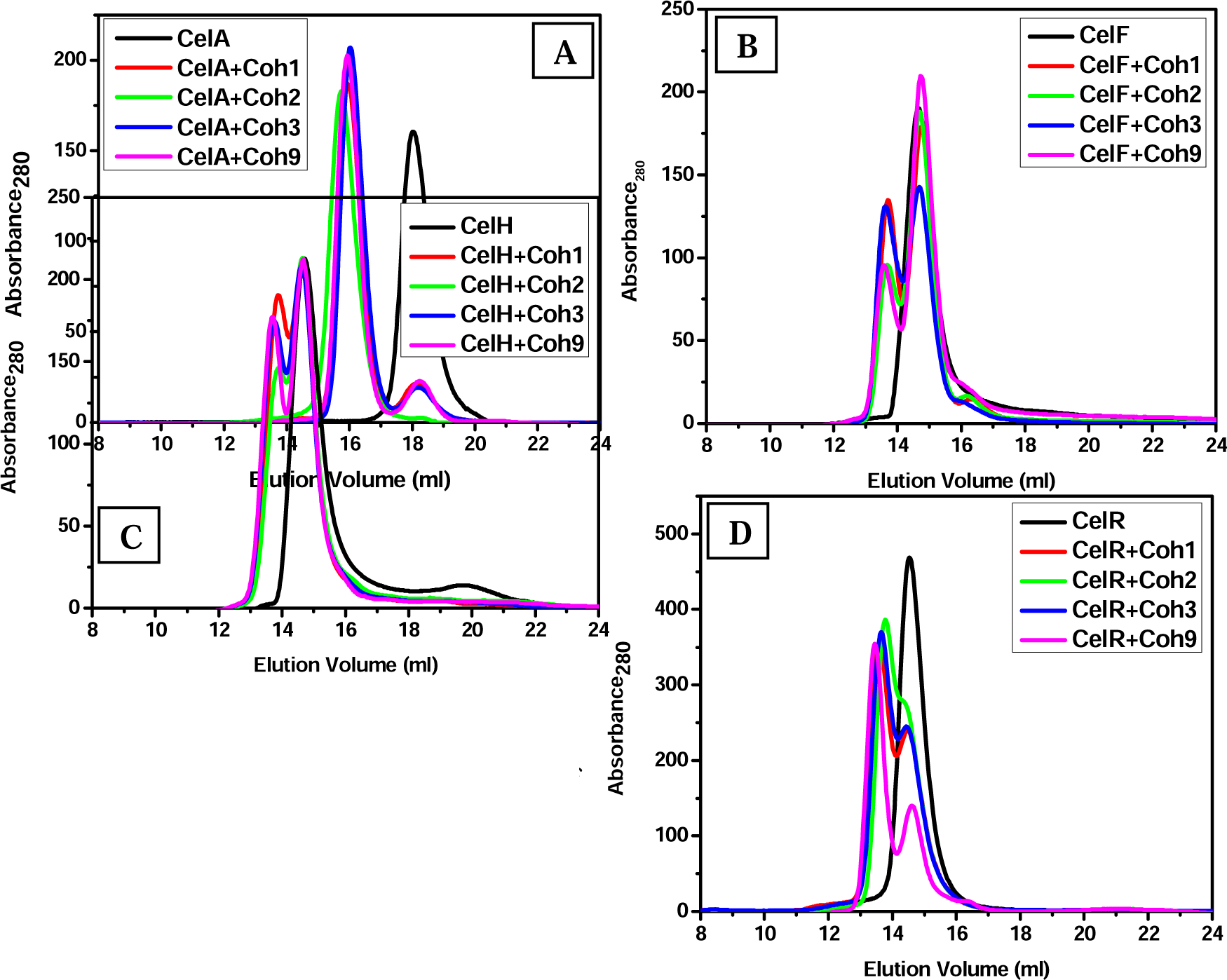
Semi-quantitative analysis of cohesin–dockerin interactions by size exclusion chromatography. Controls are shown in black color. (A) CelA (B) CelF (C) CelH and (D) CelR.

**Figure 5:**
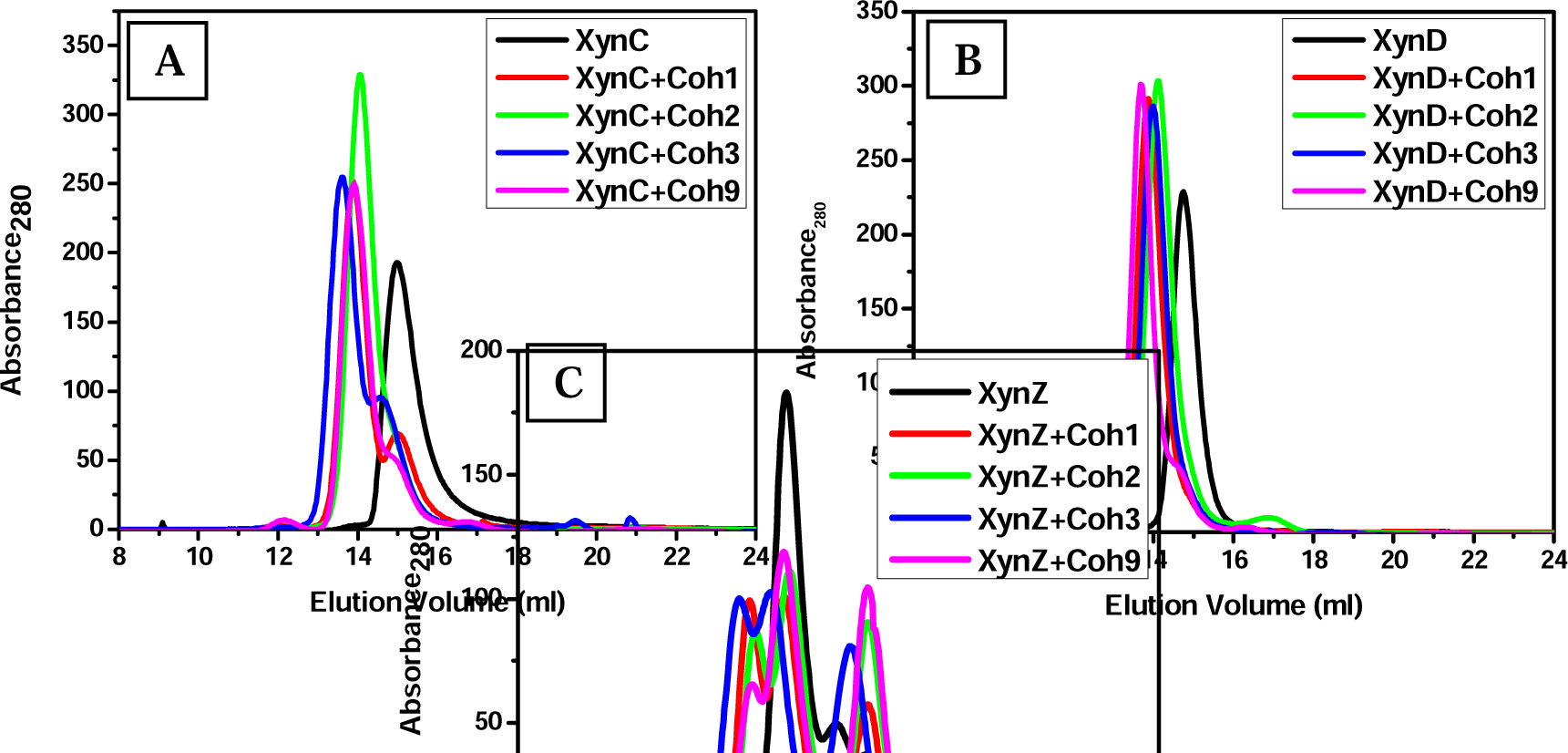
Semi-quantitative analysis of cohesin–dockerin interactions by size exclusion chromatography. Controls are shown in black color. (A) XynC (B) XynD and (C) XynZ.

**Table 2:** Comparison of preferences of cohesins derived semi-quantitatively (by SEC) and quantitatively (by PRODIGY and MST).

|  | **PRODIGY (according to pLDDT, pTM and ipTM scores)** | **SEC (according to the area under the peaks observed)** | **MST (according to estimated Kd values)** |
| --- | --- | --- | --- |
| **CelA** | Coh3 > Coh1 > Coh2 > Coh9 | Coh2 > Coh1 = Coh3 = Coh9 | Coh3 > Coh2 > Coh1 > Coh9 |
| **CelF** | Coh1 > Coh9 > Coh2 > Coh3 | Coh3 > Coh1 > Coh2 > Coh9 | Coh2 > Coh9 > Coh1 > Coh3 |
| **CelH** | Coh2 > Coh1 > Coh3 > Coh9 | Coh3 > Coh1 = Coh9 > Coh2 | Coh2 > Coh1 > Coh9 > Coh3 |
| **CelR** | Coh1 > Coh3 > Coh2 > Coh9 | Coh9 > Coh3 = Coh1 > Coh2 | Coh3 > Coh2 > Coh1 > Coh9 |
| **XynC** | Coh1 > Coh3 > Coh2 > Coh9 | Coh2 > Coh9 > Coh1 > Coh3 | Coh2 > Coh9 > Coh1 > Coh3 |
| **XynD** | Coh3 > Coh2 > Coh1 > Coh9 | Coh1 = Coh2 = Coh3 = Coh9 | Coh1 > Coh3 > Coh9 > Coh2 |
| **XynZ** | Coh3 > Coh1 > Coh2 > Coh9 | Coh1 = Coh3 > Coh2 > Coh9 | Coh1 > Coh2 > Coh3 > Coh9 |

With the cellulases, it was observed that Coh3 is preferred in two out of four cellulases. CelA would appear to show a preference for Coh2 but equal preferences for the remaining Coh domains. CelF and CelH would appear to show a preference for Coh3, but low preferences for Coh9 and Coh2. CelR shows maximum preference for Coh9 over other Coh domains. With the xylanases, XynC shows preferences for specific Coh domains, whereas XynD and XynZ appear not to prefer specific Coh domains. Thus, XynC appears to prefer Coh2, with hardly any preference for Coh3, while XynD shows equal preference for all Coh domains, leaving no unbound population of any Coh domain. In the case of XynZ, Coh1 and Coh3 were observed to be equally preferred, with Coh9 preferred to a lower extent. Overall, with the cellulases, Coh3 would seem to be the preferred Coh domain, whereas with the xylanases, at least with XynC, there is a preference for Coh2.

It is satisfying to note that although the SEC data was not entirely along the lines of the MST data, two overall conclusions were the same using data from both types of studies, namely that (i) the cellulases used for these studies all possess Doc domains that prefer to bind to Coh3, and (ii) at least one xylanase possesses a Doc domain with a clear preference for Coh2. Speaking loosely, from an evolutionary viewpoint, one could probably say that since Coh3 is nearly identical to Coh4, Coh5, Coh6, Coh7 and Coh8, and since the Doc domains of the cellulases could be assumed to have co-evolved separately from the Doc domains of the xylanases, the first two Coh domains of the CipA chain (Coh1 and Coh2) preferentially bind to xylanase-bearing Doc domains, whereas the next six Coh domains on the CipA chain (Coh3, Coh4, Coh5, Coh6, Coh7 and Coh8) all preferentially bind to cellulase-bearing Doc domains.

### Qualitative verification of Coh-Doc interactions through native PAGE (NPGE) and mass spectrometric (MS) analyses

We also verified the occurrence of all twenty-eight pairwise interactions between the four cohesin domains, and the seven enzyme-bearing Doc domains, through visualization of protein complexes as ‘emergent’ additional bands on native PAGE, in addition to bands corresponding to Coh and enzyme-bearing Doc domains. As seen in Figure 6, such emergent bands were observed as a band (or bands) of higher or lower mobility in comparison with the band corresponding to the enzyme-bearing Doc domain, since native PAGE separates complexes on the basis of both charge and size and it is conceivable for a complex to be characterized by a charged state that causes the complex to display either trend of mobility with respect to the larger-sized partner in the complex (the enzyme-bearing Doc domain), where the smaller-sized partner (the Coh domain) is considerably smaller. Prior to electrophoresis on native PAGE, enzyme-bearing Doc domains and Coh domains were mixed in equimolar concentrations of 2.5 µM, and incubated overnight. The native PAGE results confirm that essentially all enzyme-bearing Doc domains bind to all Coh domains, i.e., that there is considerable non-specificity of interactions as might be anticipated. In each panel in Figure 6, corresponding to interactions of a single enzyme-bearing Doc domain with all four Coh domains, lanes 6 through 9 display emergent bands for interactions with Coh1, Coh2, Coh3 and Coh9, respectively. We also observe some degradation, leading to enzymes that have lost the status of being in fusion with the Doc domain. Thus, in each panel, the lowest band in the control lane for the enzyme alone (corresponding to enzyme detached from its Doc domain), altered mobility is not observed. Otherwise, prominent bands are observed, which indicate the occurrence of interactions that are stable to native PAGE analyses. We did not analyse the intensities of these bands any further, or attempt to compare this qualitative evidence with results from SEC analyses, because densitometric analyses of gel data is of poorer reliability for quantitative inferences than the data from the SEC or MST analyses. However, we did subject the emergent bands (corresponding to complexes) to peptide mass fingerprinting using MALDI-MS analyses, for two representative (pairwise) interactions. As shown in Supplementary Figure 17, we were able to verify the presence of tryptic peptides from both interaction partners, i.e., the Coh domain and the enzyme-bearing Doc domain, with both emergent bands.

**Figure 6:**
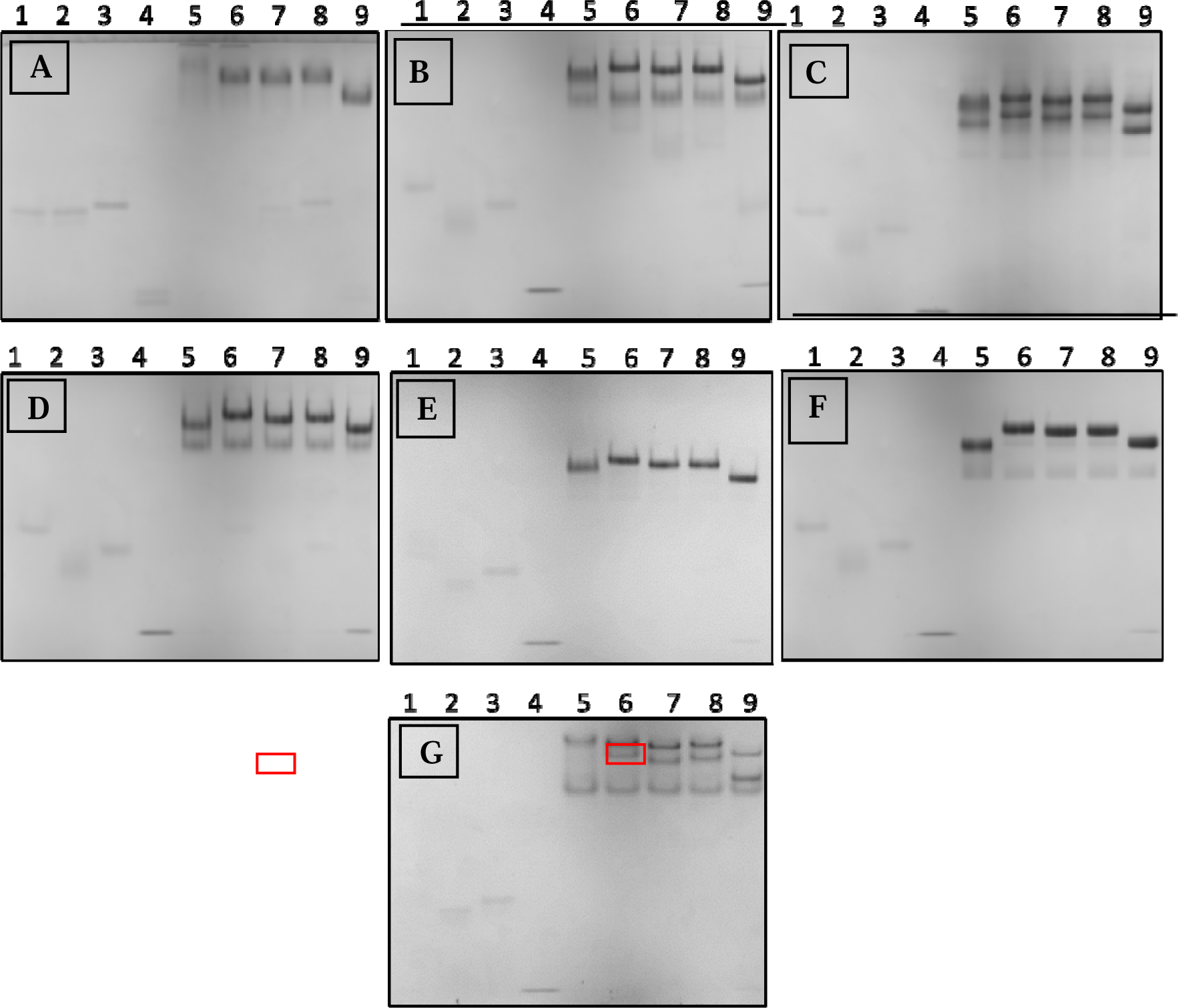
Qualitative analysis of cohesin–dockerin interactions by native PAGE. (A) CelA (B) CelF (C) CelH (D) CelR (E) XynC (F) XynD (G) XynZ. Lane numbers: (1) Coh1 (2) Coh2 (3) Coh3 (4) Coh9 (5) Enzyme (6) Enzyme+Coh1 (7) Enzyme+Coh2 (8) Enzyme+Coh3 (9) Enzyme+Coh9. Bands shown in the red boxes were selected for identity confirmation by PMF using MALDI-Q-TOF analyses on a Synapt G2S-HDMS mass spectrometer.

### Computational (quantitative) bioinformatics-based interaction (BIBA) analyses

In order to first examine general evidence for two binding modes of Coh domains to Doc domains in the structural databases, we examined two known structures (PDB IDs 1OHZ, and 4FL4). It must be noted that these structures correspond to two different Coh domains (Coh2 and Coh9) and also two different Doc domains (derived from *C. thermocellum* enzymes not used in the present study). Therefore, these structures shed no specific light on the possibility of there being two modes of binding of the same Doc domain to the same Coh domain, using helix 1 or helix 3 on the Doc domain to mediate two alternative modes of binding. Despite this, and because Doc domains share significant sequence homology, as do Coh domains, we think that each Doc domain could be imagined to be a ‘proxy’ for every other Doc domain, and that each Coh domain could likewise be imagined to be a proxy for every other Coh domain of the same type (i.e., type I cohesin), at least in respect of the overall fold, although not in respect of the microstructural features of the surfaces of helices 1 and 3. A sequence alignment of Doc domains from 1OHZ, and 4FL4, is shown in Supplementary Figure S18. The structural alignment between the Coh-Doc complexes of 1OHZ, and 4FL4, is shown in Figure 7. From this figure, it is clearly evident that Coh2 in 1OHZ (in red), and Coh9 in 4FL4 (in blue), are structurally superimposable to a significant degree, despite showing a pairwise percent sequence identity of only ∼67.14 %. The partner Doc domains (in orange and green), which have a percent sequence identity of 56.52 %, however, are not aligned, because binding involves helix 1 of the Doc domain of 1OHZ with Coh2, but helix 3 of the Doc domain of 4FL4 with Coh9. In both interactions, however, the conserved Ser/Thr pairs, e.g., Ser10/Thr11 in helix 1, and Ser46/Ser47 in helix 3, are involved in the binding to the partner Coh domain. In the two structures, the dockerin is rotated by 180 ° between one structure and the other, with respect to the Coh domain, to allow helix 3 to bind in place of helix 1.

**Figure 7:**
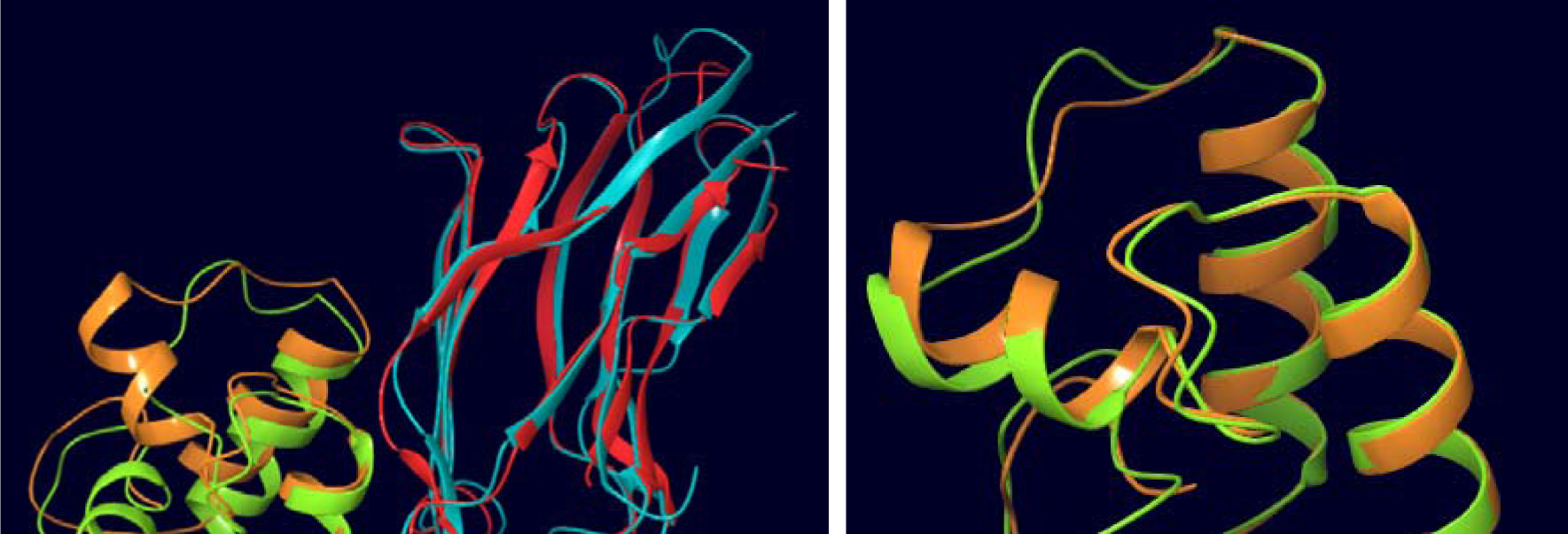
Structures are aligned in presence and absence of cohesin chains. Red and orange represents Coh2 and its dockerin partner, respectively (PDB ID: 1OHZ); Blue and green represents Coh9 and its dockerin partner, respectively (PDB ID: 4FL4).

Structure predictions for Coh domains and Doc domains (note: without enzyme in fusion with the Doc domain) were done using the Colab Fold (AlphaFold2 multimer) web server. Based on pLDDT, pTM and ipTM scores, models taken from the output from Colab Fold were ranked. Supplementary Figure 19 shows the workflow diagram used for computational analyses of pairwise interactions amongst Coh and Doc domains. For each of the 28 pairs of Coh-Doc interactions, the top 10 predicted structures were selected and examined. For 26 pairwise interactions, these included at least one case each of binding modes involving helices 1 or 3, and the top structure for each mode was selected. Interaction analyses were then carried out to predict K_d_ values for these pairs of Coh-Doc interactions at a simulated temperature of 25 L, using PRODIGY web server. For the remaining two pairwise Coh-Doc interactions (Coh2 with the Doc domain of XynZ, and Coh9 with the Doc domain of CelA), only a single mode of interaction was observed amongst all 10 top structures predicted, and this mode was subjected to interaction strength analyses. Table 3 shows the obtained K_d_ values, while Supplementary Table ST1 compares them with the experimental results from MST and SEC analyses. Notably, the bulk of pairwise interactions were predicted to occur with K_d_ values in the nanomolar (or tens of nanomolar) range. Figure 8 shows K_d_ comparisons between the two predicted binding modes. Interestingly, amongst all cohesins, Coh1 shows the strongest interactions with Doc domains, with the lowest predicted K_d_ values for binding involving both helices 1 (binding mode 1) and 3 (binding mode 2). In most cases, binding mode 2 was predicted to be stronger than binding mode 1. With Coh3 and Coh9, the difference in predicted K_d_ values for binding involving the two modes was the largest. Similarly, amongst dockerin-bearing enzymes, the differences in predicted K_d_ values for binding involving the same binding mode, to different cohesins, was high for CelH, CelR and XynC.

**Figure 8:**
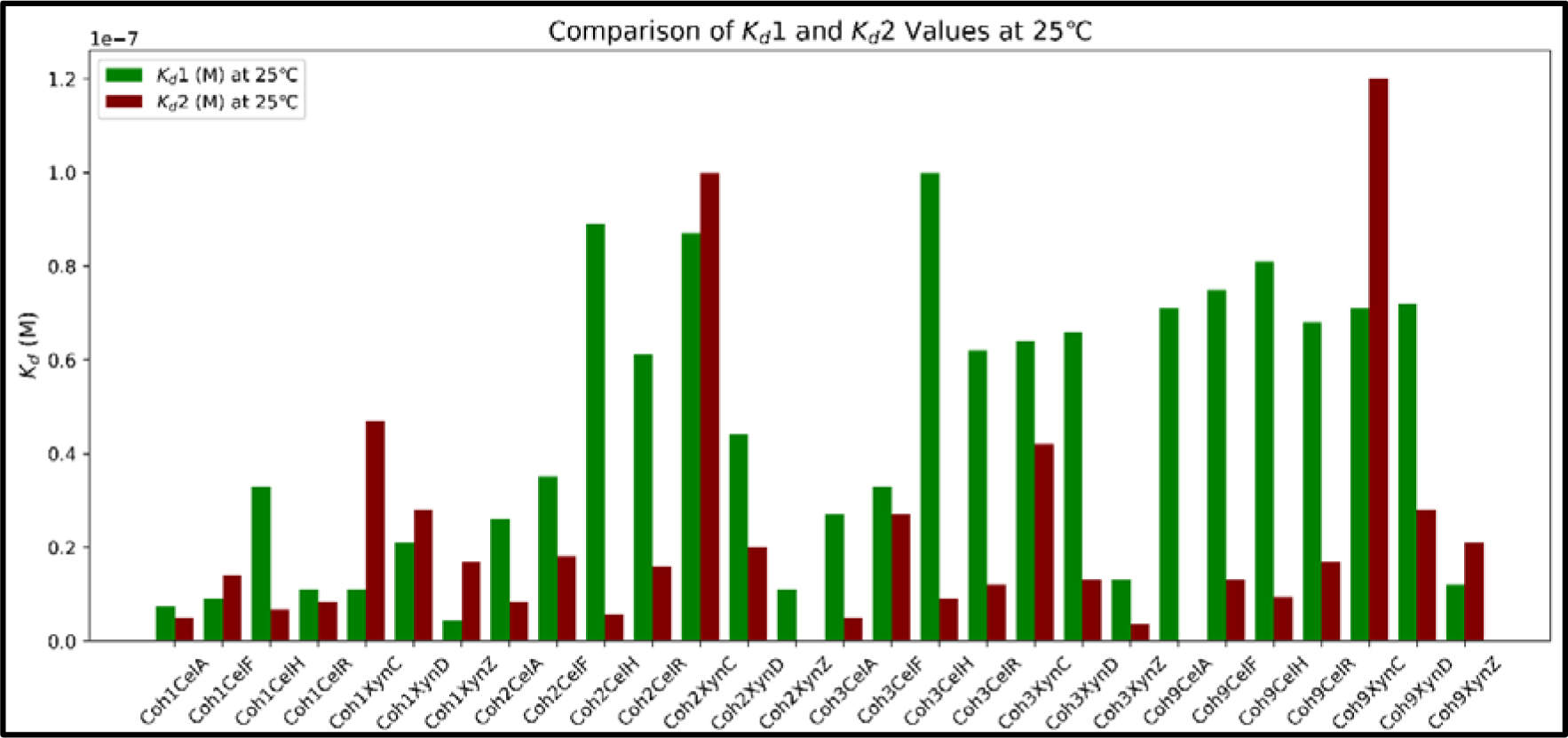
K_d_s comparison of binding mode 1 and binding mode 2 calculated using PRODIGY. Binding mode 1 (green bars) and binding mode 2 (red bars) are represented with blue and orange bars respectively.

**Table 3:** K_d_s calculated using PRODIGY for two modes of binding in all 28 pairs of cohesins and dockerins at 25 °C.

|  | Coh1 | Coh2 | Coh3 | Coh9 |
| --- | --- | --- | --- | --- |
| **CelA** | 7.3 nM and 4.9 nM | 26 nM and 8.4 nM | 27 nM and 5 nM | 71 nM |
| **CelF** | 9 nM and 14 nM | 35 nM and 18 nM | 33 nM and 27 nM | 75 nM and 13 nM |
| **CelH** | 33 nM and 6.6 nM | 89 nM and 5.8 nM | 100 nM and 9 nM | 81 nM and 9.4 nM |
| **CelR** | 11 nM and 8.4 nM | 61 nM and 16 nM | 62 nM and 12 nM | 68 nM and 17 nM |
| **XynC** | 11 nM and 47 nM | 87 nM and 100 nM | 64 nM and 42 nM | 71 nM and 120 nM |
| **XynD** | 21 nM and 28 nM | 44 nM and 20 nM | 66 nM and 13 nM | 72 nM and 28 nM |
| **XynZ** | 4.3 nM and 17 nM | 11 nM | 13 nM and 3.6 nM | 12 nM and 21 nM |

## Discussion

In this paper, we have presented a detailed, multi-faceted, and multi-pronged analysis of domain-domain interactions in *C. thermocellum* cellulosomes. Our goal was to understand why there are multiple cohesin (Coh) and dockerin (Doc) domains with different amino acid sequences, but similar overall three-dimensional folds. We performed an exploration of interactional preferences amongst different chosen Coh and Doc domains, to ask whether such preferences could potentially play a role in determining the likely location of particular enzyme-bearing Doc domains upon the CipA (scaffoldin) polypeptide chain containing nine Coh domains. We did this by producing recombinant protein forms of four representative Coh domains and seven enzyme-bearing Doc domains. We then studied pairwise interactions amongst Coh and Doc domains, using quantitative data from microscale thermophoresis experiments, and semi-quantitative data from size exclusion chromatography experiments. We have established conclusively that: (i) all Coh domains interact with all Doc domains, amongst representatives studied; (ii) some pairs of Coh and Doc domains display two modes of binding (potentially akin to the dual modes of binding earlier noted in crystal structures of different pairs of interacting Coh and Doc domains), using different affinities for the two modes; (iii) interactional preferences of xylanase-associated Doc domains exist for two Coh domains (Coh1 and Coh2); (iv) interactional preferences of cellulase-associated Doc domains exist for Coh3, which is virtually identical to five other domains (Coh4-to-Coh8) in CipA; (v) Coh9 has high affinities for the Doc domains of a xylanase, XynC, and a cellulase, CelR, but poor affinities for all other partners tested. Overall, our data suggests that if everything else is equal, and if all enzyme-bearing Doc domains are present at equimolar concentrations, xylanase and cellulase enzymes, respectively, could potentially locate differentially upon CipA chain, with the former locating upon the two N-terminal domains, and the latter upon the broad middle of the chain, with Coh9 at the C-terminus binding to specific xylanase or cellulase partners. An extrapolation of our data would also suggest that steric adjustments between large enzymes that attempt to locate upon adjacent Coh domains on CipA could be resolved through use of alternative modes of binding of some enzyme-bearing Doc domains.

Additionally, qualitative data from native gel electrophoresis experiments is presented in support of the belief that all Coh domains are capable of binding to all Doc domains. Also, data from computational studies is presented in support of the contention that the majority of Coh-Doc interactions can potentially occur in either of the two alternative binding modes, albeit with different affinities. An interesting and curious aspect of our work is the finding (from both quantitative microscale thermophoresis experiments, and semi-quantitative size exclusion chromatography experiments) that Coh9 binds weakly to most, but not all, enzyme-bearing Doc domains tested. This suggests that Coh9, which is located near the C-terminus of CipA (just prior to the terminal type II dockerin domain), tends to be occupied only by enzymes present at high enough concentrations to ensure its occupancy. Our bioinformatics-based analyses also point towards Coh9 displaying the weakest binding to most Doc domains associated with enzymes. We believe that if such data concerning interactional preferences were to be comprehensively collected, in pairwise fashion, for all conceivable (∼over 70) enzyme-bearing Doc domains, and for all nine Coh domains present upon CipA (including domains Coh4, Coh5, Coh6, Coh7 and Coh8, which were not tested because of their high sequence identity/homology with Coh3), it would then become conceivable for this knowledge to then be employed to produce recombinant, chimeric cellulosomes for multiple and variegated uses. It could also become possible to improve degradation of specific types of biomass by ensuring the proximity of chosen enzyme components to give rise to enhanced and synergistic degradation. Further, this understanding of Coh-Doc interactions could help in the creation of other non-cellulosomal applications that exploit the association of protein constructs incorporating Coh/Doc domains. One such instance has been recently attempted successfully by us [38].

## Materials and Methods

### Choice of Doc domain-bearing enzymes and Coh domains

A total of seven enzymes, consisting of 4 cellulases, Cel8A (also referred to as CelA), Cel9F (also referred to as CelF), Man26/5H (also referred to as CelH), and Cel9R (also referred to as CelR), and 3 xylanases, Xyn10C (also referred to as Xyn10C), Xyn11D (also referred to as XynD), and Xyn10Z (also referred to as XynZ) were used for this study, from amongst the over ∼70 Doc domain-bearing enzymes encoded by the *Clostridium thermocellum* genome. Further, from amongst the nine Coh domains present in the CipA cellulosomal chain of *C. thermocellum*, the 4 that are most dissimilar in sequence, i.e., Coh1, Coh2, Coh3, and Coh9, were chosen for this study. It may be noted that Coh1, Coh2, and Coh9, show 69, 81, and 75 % identity, respectively, with Coh3 (which shows over 95 % identity with Coh4, Coh5, Coh6, Coh7 and Coh8).

### Cloning of genes encoding Coh domains and enzyme-bearing Doc domains

All Coh domains and enzyme-bearing Doc domains were amplified from *C. thermocellum* genomic DNA (strain ATCC 27405) through PCR involving appropriately chosen primers incorporating the restriction sites, NdeI, or NheI (in the forward primer) and XhoI (in the reverse primer). The PCR products obtained were then digested by NdeI, and XhoI, in case of all cohesins, Xyn10C (or XynC) and Cel8A (or CelA), but by NheI and XhoI in case of Xyn11D (XynD), Xyn10Z (or XynZ), Cel9F (or CelF), Man26/5H (or CelH) and Cel9R (or CelR), and ligated into the T7-based expression vector, pET23a, for production in fusion with a C-terminal 6xHis affinity tag in *Esherichia coli* strain BL21 Star (DE3) pLysS, as described below.

### Expression and purification of proteins

Plasmid vectors (pET23a) bearing genes encoding the desired constructs were transformed into XL-1 Blue *E. coli* cells to produce plasmids for DNA sequencing. After sequencing-based confirmation of identity, these plasmids were transformed into *E. coli* BL21 Star (DE3) pLysS for protein expression and purification. Overexpression was induced by 1 mM IPTG in the mid-exponential phase of culture growth, at an optical density (O.D_600_) of 0.6, during growth of transformed cells in LB media. Following induction, cells were incubated for 12-16 hours at 25 °C, sedimented through centrifugation, and lysed. Expressed 6xHis-tagged protein fusions were then chromatographically purified from clarified lysates, using immobilized metal (Ni-NTA) affinity chromatography upon columns sourced from GE Healthcare. Protein yields were typically in the range of 2-3 mg per litre of bacterial culture. All proteins were purified from lysates under non-denaturing conditions, prior to transfer into 20 mM Na-HEPES buffer (pH 7.5, containing 100 mM NaCl, and 2 mM CaCl_2_). This buffer transfer was carried out through buffer-exchange using a Superdex-75 size exclusion chromatography (SEC) column (GE Healthcare) for individual Coh domains, and a Superdex-200 SEC column (GE Healthcare) for enzyme-bearing Doc domains, using a GE Akta Purifier 10 chromatographic workstation.

### Spectroscopic characterization of folded state

The secondary structures of all Coh domains and enzyme-bearing Doc domains were examined using circular dichroism, with spectra collected upon a Chirascan™ spectrometer (Applied Photophysics Ltd.), using a quartz cuvette of 1 mm path length, and a protein concentration of 0.2 mg/ml, in 20 mM HEPES buffer of pH 7.5, containing 100 mM NaCl and 2 mM CaCl_2_. Mean residue ellipticity (MRE) was calculated using the formula, MRE = (θ × MRW × 100) / (1000 × concentration in mg/ml × pathlength in cm), where 0 was the raw ellipticity measured in millidegrees, and the MRW was the exact mean residue weight calculated for the protein. Tertiary structure formation in all proteins was examined through assessment of folded state based on fluorescence spectroscopic examination of the wavelength of maximal fluorescence emission of intrinsic fluorescence, derived from tryptophan (15 to 20 tryptophan residues number in each of the 7 enzymes, with peak excitation at 282 nm and peak emission in the range of 335 to 345 nm). The occurrence of a wavelength of maximal emission significantly lower than ∼353 nm was taken to be evidence of the shielding of tryptophan residues from aqueous solvent through formation of a well-folded tertiary structure. The Coh domains contain no tryptophan residues, but at least four tyrosine residues each. Protein concentration estimation was done using the absorbance at 280 nm predicted by the Protparam module in the software, Expasy.

### Spectroscopic and calorimetric characterization of protein thermal stability

Different methods were used to assess the high thermal stability of the folded states of all Coh domains and enzyme-bearing Doc domains (anticipated due to their being derived from the genome of a thermophile, *C. thermocellum*). Firstly, differential scanning calorimetry (DSC) was used to examine thermal denaturation through heating from 20 °C to 90 °C at a rate of 60 °C/h (to assess the T_m_, or melting temperature), followed by cooling from 90 °C to 20 °C, at a rate of 60 °C/h (to assess whether refolding occurs from a thermally-unfolded state) with concomitant measurement of enthalpic changes, using a VP-DSC (MicroCal) differential scanning calorimeter, and a protein concentration of 0.5 mg/ml. DSC data was collected after 10 cycles of heating and cooling of the Na-HEPES buffer, to generate the baseline. Data concerning the heat required to unfold the protein was extracted from the system and measured, over the temperature range in which a difference in the rate of change of temperature between sample and control (reference) cells could be observed. A specific heat capacity versus temperature graph was obtained, and the area under the concentration-normalized (baseline-subtracted) graphs was estimated to measure the change in enthalpy associated with protein unfolding, with the peak of the transition curve assessed to be the measured melting temperature (T_m_). Further, thermal stability was also assessed using fluorescence spectroscopy accompanying the thermal denaturation of all proteins (during heating from 20 °C to 90 °C) to monitor the shifting of the protein’s wavelength of maximal fluorescence emission towards longer wavelengths, with changes in tertiary structure accompanying unfolding, on a steady-state FluoroMax (Horiba) fluorimeter with excitation at 295 nm, and emission recorded between 300 and 400 nm. Later, the ratio of emissions at 330 and 350 nm was plotted.

### Analyses of Coh-Doc interactions by microscale thermophoresis (MST)

Microscale thermophoresis was performed to study Coh-Doc interactions, using Coh domains and Doc domain-bearing enzymes. The purified Coh domains were labelled using red-NHS 2^nd^ generation dye, according to the manufacturer’s instructions. Binding assays were performed with a Monolith NT.115 device (a trademark of NanoTemper Technologies GmbH, Munich, Germany) using standard treated capillaries. To improve the accuracy of the K_d_ determination, the concentration of labelled Coh protein was restricted to 100 nM, yielding a fluorescence signal strength of above 200 units. Equal concentrations of labelled protein (100 nM) were titrated against varying ligand (protein) concentrations (10 µM to 0.06 nM) of the enzyme-bearing Doc domains, with change in distribution of fluorescence from the Coh domains upon heating (by 2-6 °C, using an IR laser) measured as a function of the concentration of the protein-protein complex constituted by Coh domains bound to enzyme-bearing Doc domains. Since migration of an individual molecule differs from migration of a molecule bound to ligand, the change in distribution of fluorescence was used to determine the ratio of free protein to protein-protein complex. F_cold_ and F_hot_ were used to measure the fluorescence before and after heating, respectively. F_hot_/F_cold_ gave the normalized fluorescence with plots F_norm_ against the logarithmic concentrations of serially diluted ligand, yielding sigmoidal binding curves.

### Analyses of Coh-Doc interactions by size exclusion chromatography (SEC)

Semi-quantitative analyses of interactions between different Coh domains and enzyme-bearing Doc domains were performed using size exclusion chromatography. Pairs of proteins (5 µM of each) were constituted in 20 mM HEPES buffer, pH 7.5, and incubated for ∼12 hours (overnight) at room temperature and loaded upon a Superdex-200 Increase 10/300 GL gel filtration column in an AKTA Purifier-10 workstation (GE Healthcare). The enzyme-bearing Doc domains were also chromatographed, in a separate run, as a control. A reduction in the elution volume of the enzyme-bearing Doc domains population (i.e., a shift to the left) due to increase in hydrodynamic volume upon complexation with a Coh domain was monitored, to verify that an interaction had occurred, through comparison with the elution of the control enzyme-bearing Doc domains population. The extent of the shift of the complexed population, as a function of residual (uncomplexed) population eluting at the original volume (for control enzyme-bearing Doc domain) was turned into a semi-quantitative assessment of complex formation.

### Analyses of Coh-Doc interactions by native polyacrylamide gel electrophoresis (NPGE)

A qualitative analysis of interactions between Coh domains and Doc domain-bearing enzymes was also performed using native PAGE. Final concentrations of 2.5 µM of each protein were incubated at room temperature overnight and loaded upon a 10 % native PAGE gel after mixing with sample loading buffer. The assembly was kept on ice and subjected to electrophoresis for 2-3 hours at 70 V. Gels were stained, washed with water, and destained to analyze band position. Controls of Coh domains and Doc domain-bearing enzymes were also subjected to electrophoresis in separate lanes. Images were documented using Gel Doc EZ system and ImageLab software (BioRad).

### Analyses of Coh-Doc properties and interactions by bioinformatics-based approaches (BIBA)

Protein structure (PDB) files were downloaded from the RCSB Protein Data Bank. Pymol (Schrodinger) was used for structural visualization. Using the Google Colab implementation of AlphaFold, the structures of all protein complexes were predicted using AlphaFold 2 Multimer (v2.x). Multiple structural predictions were generated for each protein-protein interaction, and the best model was chosen for each binding mode based on visual inspection of key interaction sites and confidence scores. The PRODIGY server (PROtein binDIng enerGY prediction) was used to estimate dissociation constants (K_d_s) for chosen pairs at 25 °C, and assess the strength of the interaction computationally, prior to experimental determination of the same, to enable comparisons. Two or more protein sequences were analyzed using Clustal Omega (EMBL-EBI). Jalview v2.11.3.3 was used to visualize sequence alignments. Sequence logo creation was done with a web application known as WebLogo 3. Consensus sequences for various *C. thermocellum* Doc and Coh domains were created using WebLogo 3.

## Supporting information

All supplementary figures and tables

## Acknowledgements

We thank the Government of India for financial support (for equipment/consumables) extended through IISER Mohali. Gurmeet Kaur is acknowledged for assistance rendered during some experiments involving assessment of interactions by size exclusion chromatography.

We thank the Ministry of Education, Government of India for a Centre of Excellence grant in Protein Science, Design, and Engineering (CPSDE; MHRD-14-0064), awarded to P.G., the Department of Biotechnology, Government of India for a grant of a Hyperthermophile Enzyme Hydrolase Research Centre (HEHRC; BT/PR/31706/PBD/26/705/2019) awarded to P.G., and the TATA Transformation Prize Grant, awarded to P.G. S.W. thanks the University Grants Commission (UGC) for a research fellowship. V.Y. thanks IISER Mohali for a merit-cum-means scholarship.

## Funding

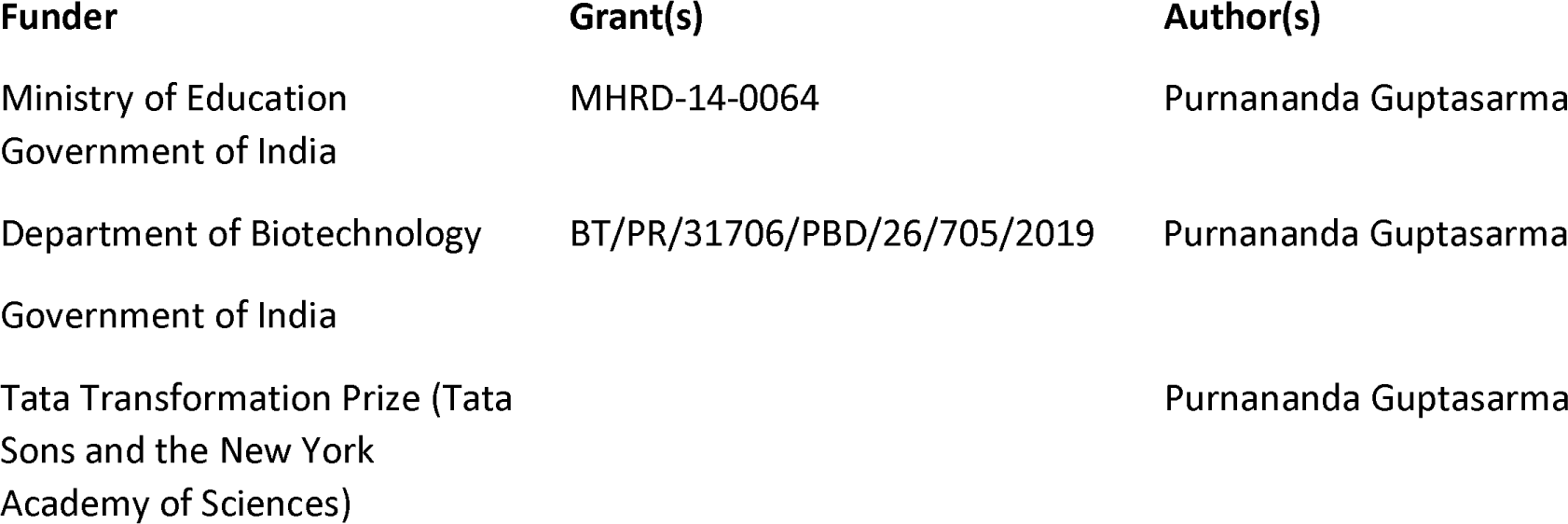

## Author Contributions

Snehal Waghmare, Conceptualization, Data curation, Formal analysis, Investigation, Methodology, Writing – original draft, Writing – review and editing | Vikas Yadav, Data curation, Formal analysis, Investigation, Methodology, Writing – original draft, Writing – review and editing | Saji Menon, Data curation, Formal analysis | Purnananda Guptasarma, Conceptualization, Formal analysis, Funding acquisition, Investigation, Methodology, Project administration, Resources, Supervision, Writing – original draft, Writing – review and editing

## Data Availability

All the data for this manuscript are contained in the article above and the supplementary information file.

## Additional Files

The following material is available online.

## Supplemental Material

Supplemental material (JB00XXX-25-s000X.docx). Fig. S1 to S19.

