## Supplementary material for "Exploring differential interactional preferences of enzyme-bearing dockerins for cohesin domains in the *Clostridium thermocellum* cellulosome": All supplementary figures and tables

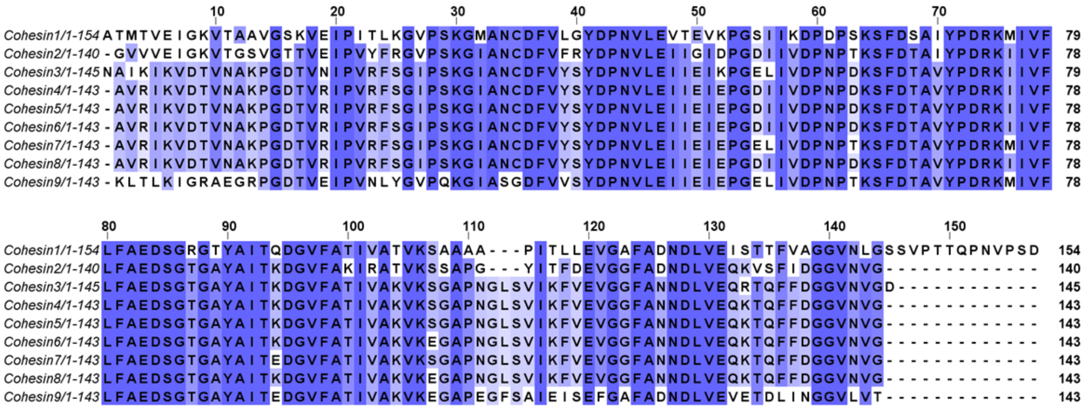


**Supplementary Figure S1:** Multiple Sequence Alignment (MSA) performed using UniProt align (Clustal Omega) showing highly conserved regions in all nine Coh domains of the polypeptide chain of CipA (Note: Color shades are proportional to percent identity, with darker shades representing residues that are more highly conserved).

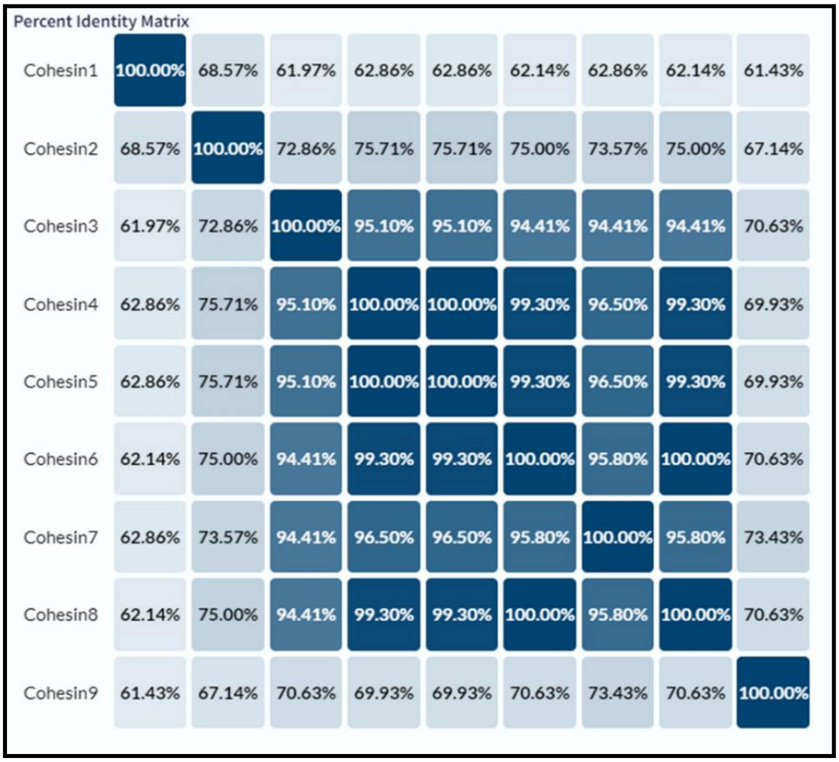

Percent Identity Matrix

| | | | | | | | | | |
|---|---|---|---|---|---|---|---|---|---|
| Cohesin1 | 100.00% | 68.57% | 61.97% | 62.86% | 62.86% | 62.14% | 62.86% | 62.14% | 61.43% |
| Cohesin2 | 68.57% | 100.00% | 72.86% | 75.71% | 75.71% | 75.00% | 73.57% | 75.00% | 67.14% |
| Cohesin3 | 61.97% | 72.86% | 100.00% | 95.10% | 95.10% | 94.41% | 94.41% | 94.41% | 70.63% |
| Cohesin4 | 62.86% | 75.71% | 95.10% | 100.00% | 100.00% | 99.30% | 96.50% | 99.30% | 69.93% |
| Cohesin5 | 62.86% | 75.71% | 95.10% | 100.00% | 100.00% | 99.30% | 96.50% | 99.30% | 69.93% |
| Cohesin6 | 62.14% | 75.00% | 94.41% | 99.30% | 99.30% | 100.00% | 95.80% | 100.00% | 70.63% |
| Cohesin7 | 62.86% | 73.57% | 94.41% | 96.50% | 96.50% | 95.80% | 100.00% | 95.80% | 73.43% |
| Cohesin8 | 62.14% | 75.00% | 94.41% | 99.30% | 99.30% | 100.00% | 95.80% | 100.00% | 70.63% |
| Cohesin9 | 61.43% | 67.14% | 70.63% | 69.93% | 69.93% | 70.63% | 73.43% | 70.63% | 100.00% |

**Supplementary Figure S2:** Percent identity matrix obtained using UniProt align (Clustal Omega) for all nine Coh domains of the polypeptide chain of CipA. Note that Coh domains Coh3 to Coh 8 share over 94% identity. Coh domains, Coh4 and Coh 5 share 100% identity, as do Coh domains, Coh 6 and 8.

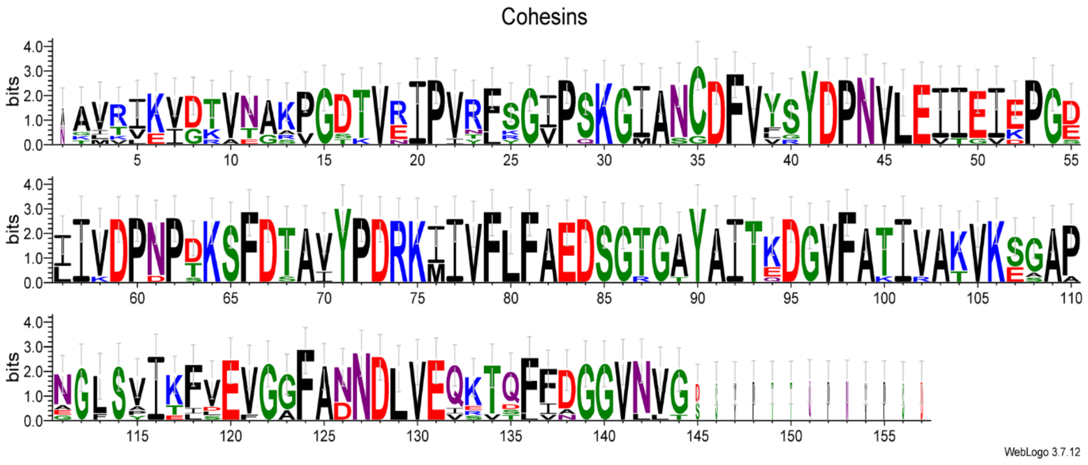


**Supplementary Figure S3**: Sequence logo representation for all nine Coh domains of the CipA polypeptide chain, showing residue conservation information and occurrence frequencies in terms of the heights of letters representing individual amino acids, generated using WebLogo 3.

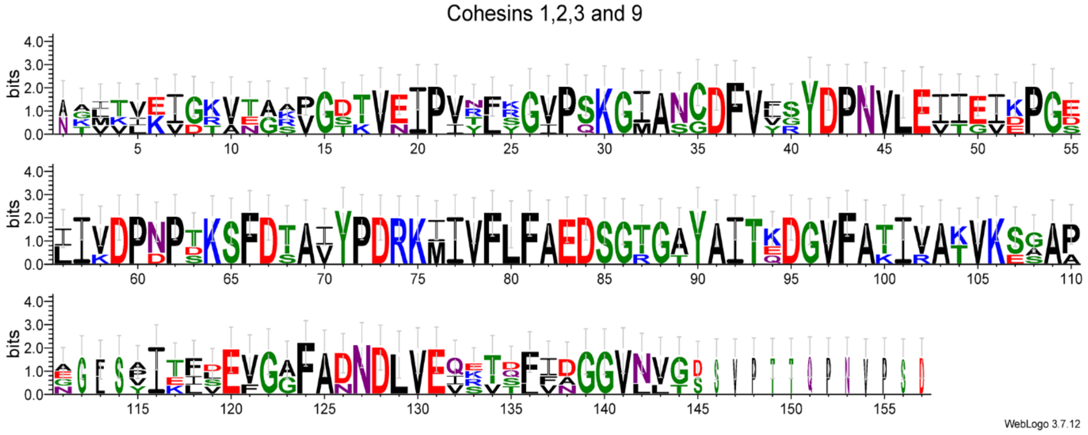


**Supplementary Figure S4**: Sequence logo representation for the four selected Coh domains, Coh1, Coh2, Coh3 and Coh9, of the polypeptide chain of CipA, showing residue conservation information and occurrence frequencies in terms of the heights of letters representing individual amino acids, generated using WebLogo 3.

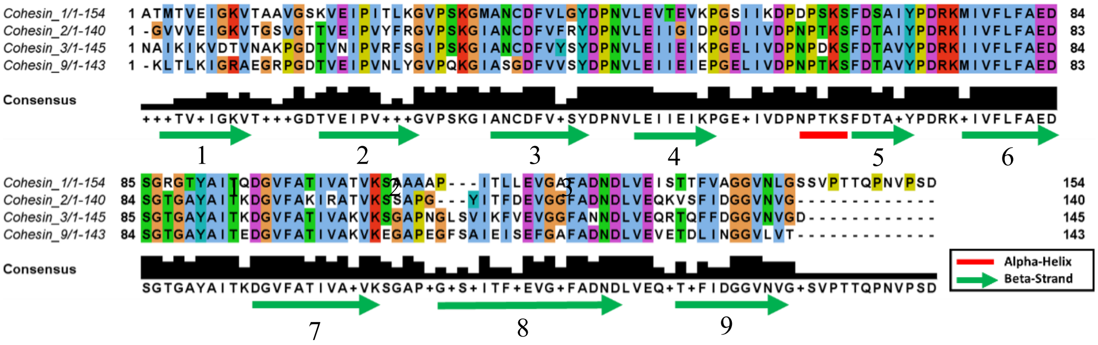


**Supplementary Figure S5**: Multiple sequence alignment showing conservation among selected Coh domains (Coh1, Coh2, Coh3 and Coh9). Clustal omega color scheme has been followed. Sequences corresponding to the secondary structural elements of these domains, based on known crystal structures of Coh2, Coh7 (which is identical to Coh3) and Coh9, are shown using green arrows (beta strand) and red lines (helix) on this multiple sequence alignment scheme. Reading from the N-terminus at the top left of the scheme, the beta strands at positions 3, 5, 6 and 8 are involved in interactions with enzyme-bearing Doc domains.

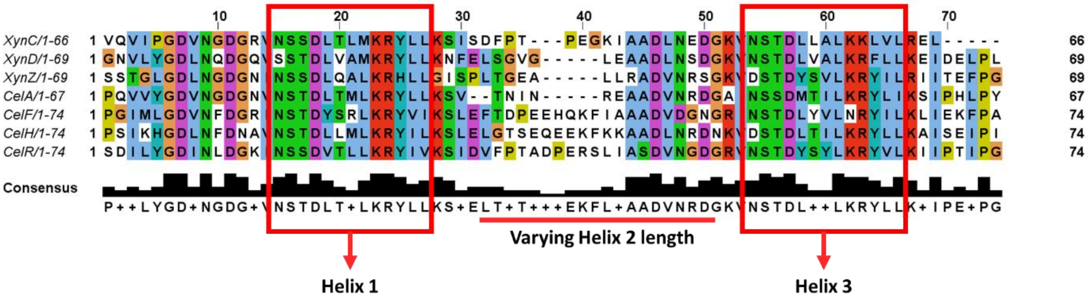


**Supplementary Figure S6**: Multiple sequence alignment scheme showing sequence conservation amongst dockerin (Doc) regions of seven selected enzymes produced and examined in this study. Clustal omega color scheme has been followed. Sequence regions corresponding to Helix 1 and Helix 3 are marked out using red boxes, and can be seen to be nearly identical. In a mutually exclusive manner, only one of these helices is responsible for interaction with the beta strands 3, 5, 6 and 8 of any cohesin (Coh) domain (see Supplementary Figure S5). The NSTD sequence at beginning of each helix contains adjacent ‘S’ and ‘S’/’T’ residues which are thought to be important for cohesin-dockerin interactions.

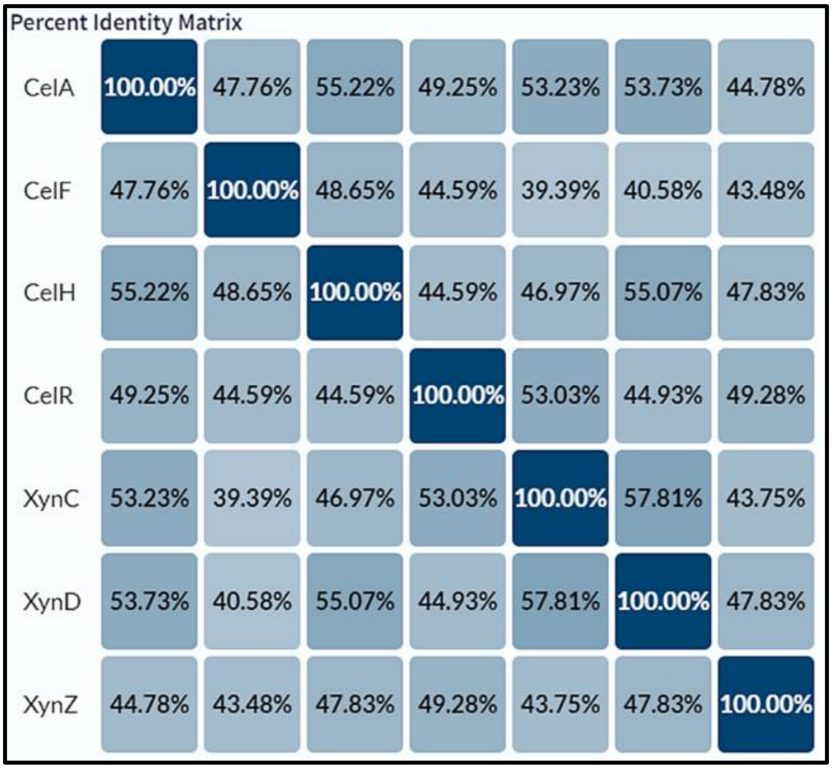
Percent Identity Matrix

| | | | | | | | |
|---|---|---|---|---|---|---|---|
| CelA | 100.00% | 47.76% | 55.22% | 49.25% | 53.23% | 53.73% | 44.78% |
| CelF | 47.76% | 100.00% | 48.65% | 44.59% | 39.39% | 40.58% | 43.48% |
| CelH | 55.22% | 48.65% | 100.00% | 44.59% | 46.97% | 55.07% | 47.83% |
| CelR | 49.25% | 44.59% | 44.59% | 100.00% | 53.03% | 44.93% | 49.28% |
| XynC | 53.23% | 39.39% | 46.97% | 53.03% | 100.00% | 57.81% | 43.75% |
| XynD | 53.73% | 40.58% | 55.07% | 44.93% | 57.81% | 100.00% | 47.83% |
| XynZ | 44.78% | 43.48% | 47.83% | 49.28% | 43.75% | 47.83% | 100.00% |

**Supplementary Figure S7**: Percent identity matrix of the sequences of the dockerin regions of the seven enzymes selected for studies, constructed using UniProt align (Clustal Omega).

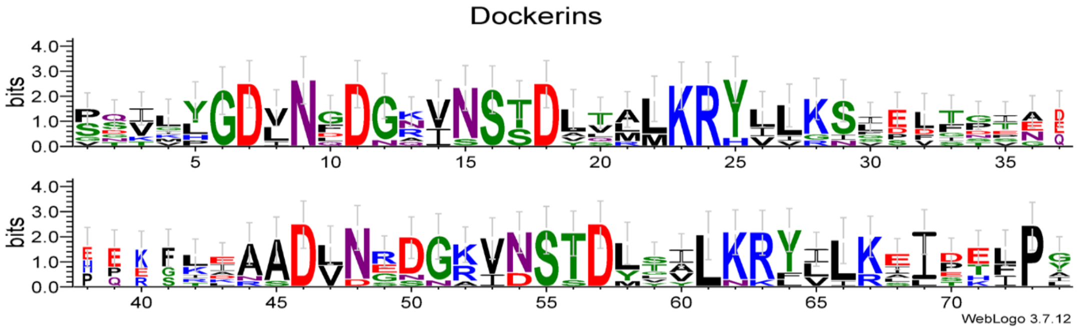


**Supplementary Figure S8**: Sequence logo representation of the Doc domains of all seven selected enzymes, showing residue conservation information and occurrence frequencies in terms of the heights of letters representing individual amino acids, generated using WebLogo 3.

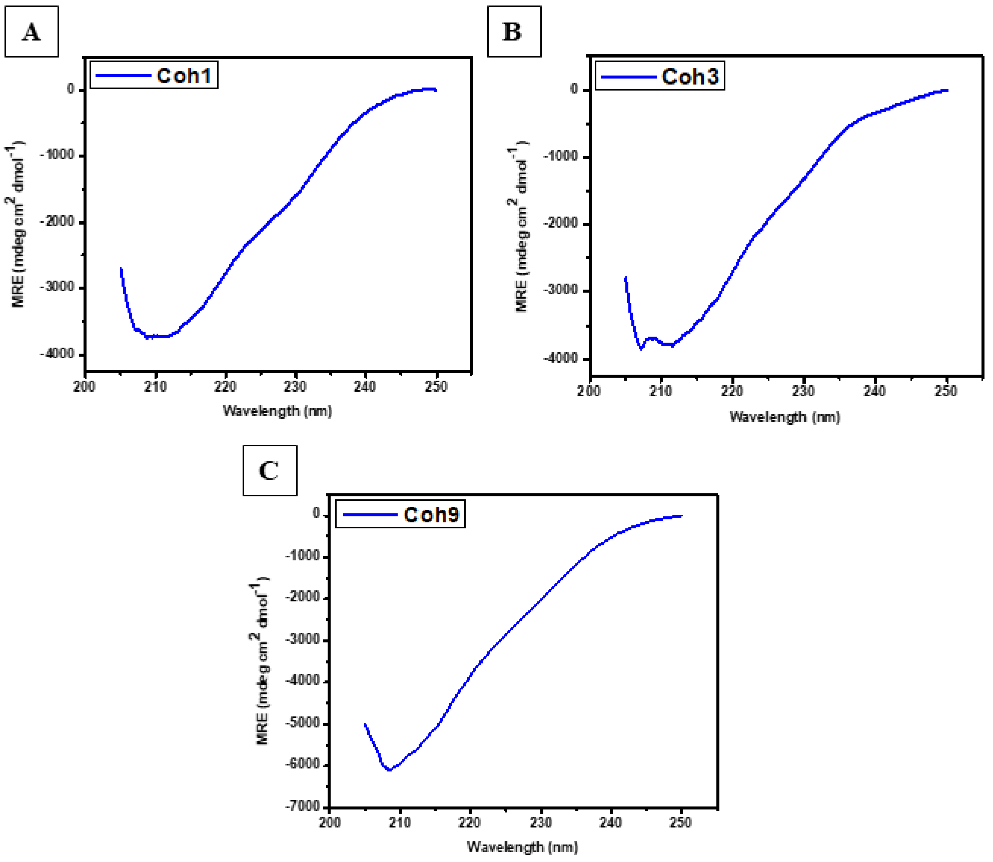


**Supplementary Figure S9**: Circular Dichroism (CD) spectra of Coh domains, Coh1, Coh3 and Coh9. The CD spectrum of Coh2 is shown in the main manuscript, Figure 2C. The CD data help to establish the folded nature of the Coh domains.

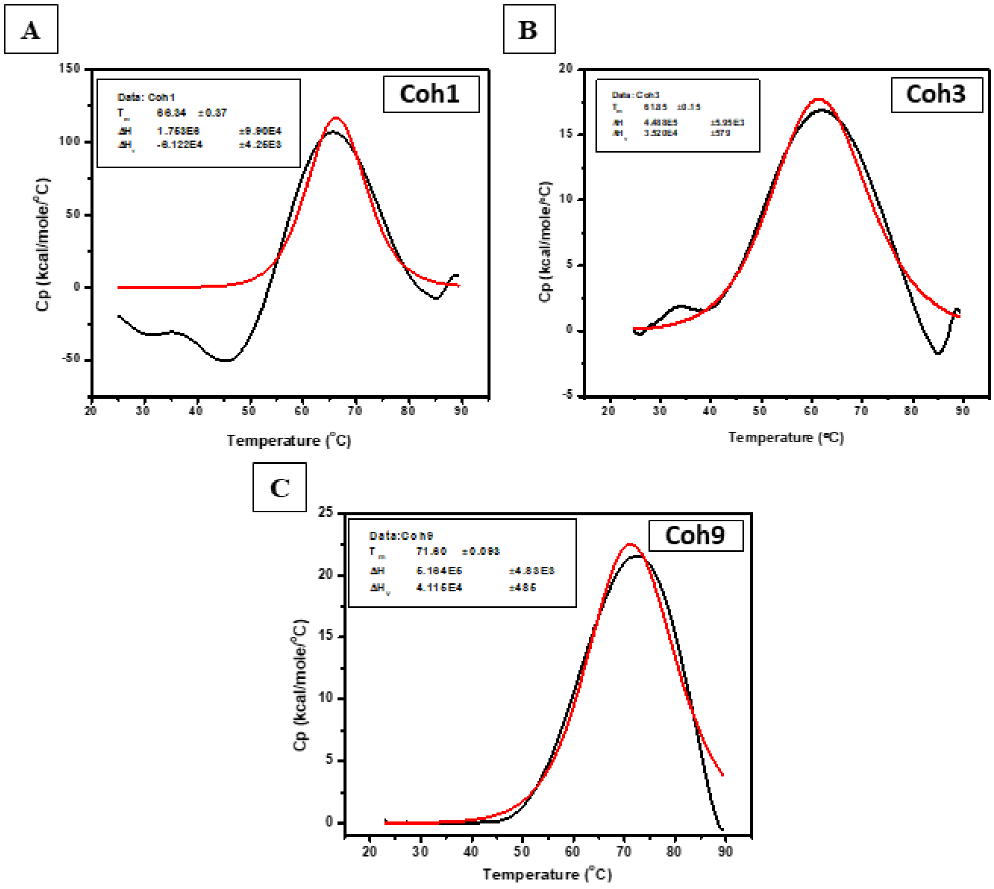


**Supplementary Figure S10**: Differential scanning calorimetry (DSC) data for the heating of Coh1, Coh3 and Coh9, with the data shown in black and the fit to a Gaussian melting curve shown in red. The data for Coh2 is shown in the main manuscript, Figure 2E. The DSC data help to establish the thermostability of the recombinant Coh domains.

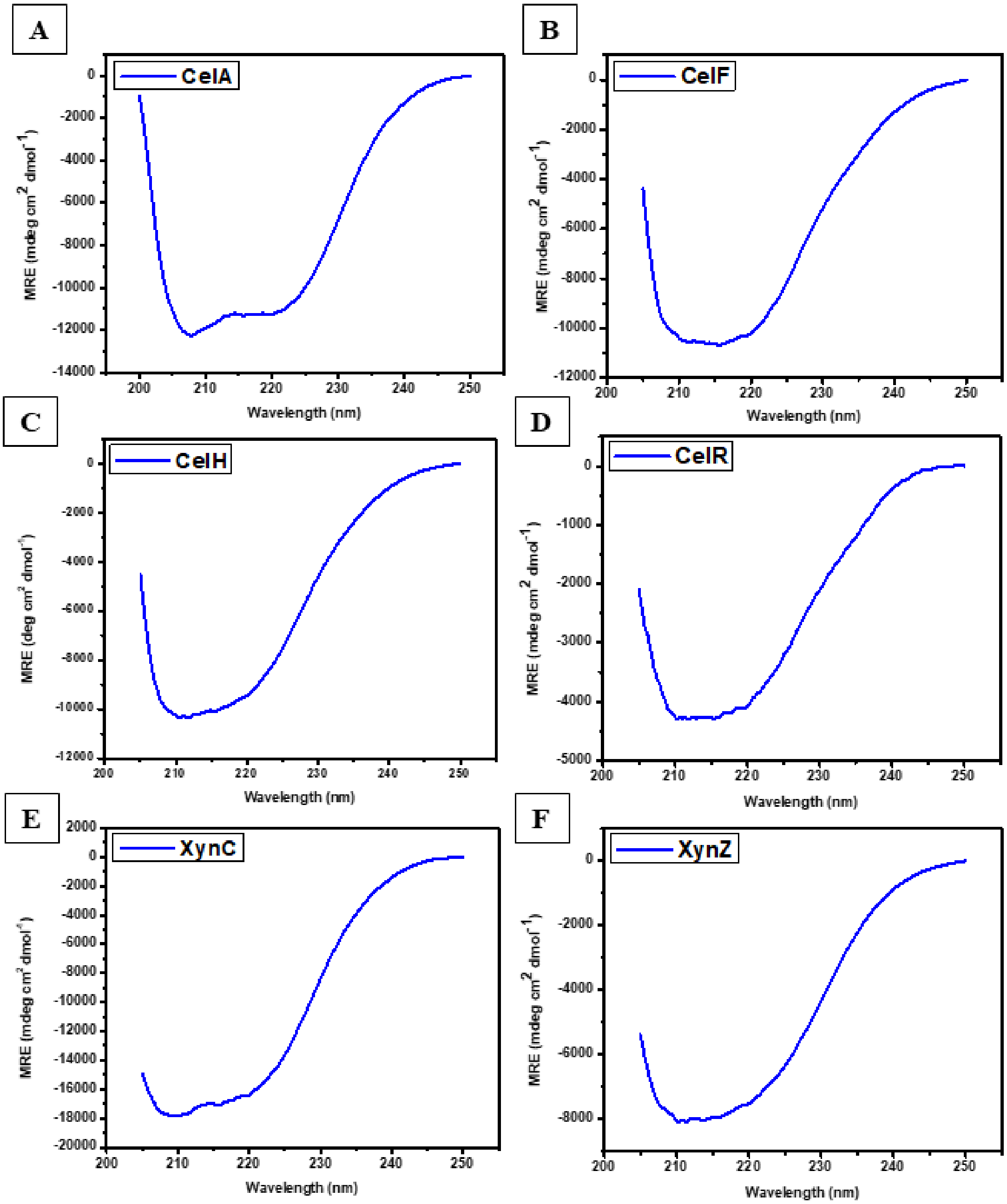


**Supplementary Figure S11**: Circular Dichroism (CD) spectra of enzyme-bearing Doc domains, shown for six of the seven enzymes selected for this study (the seventh is shown in the main manuscript, Figure 2D. The CD spectra help to establish the folded natures of the enzymes and also their relative overall contents of alpha helical and beta sheet structures.

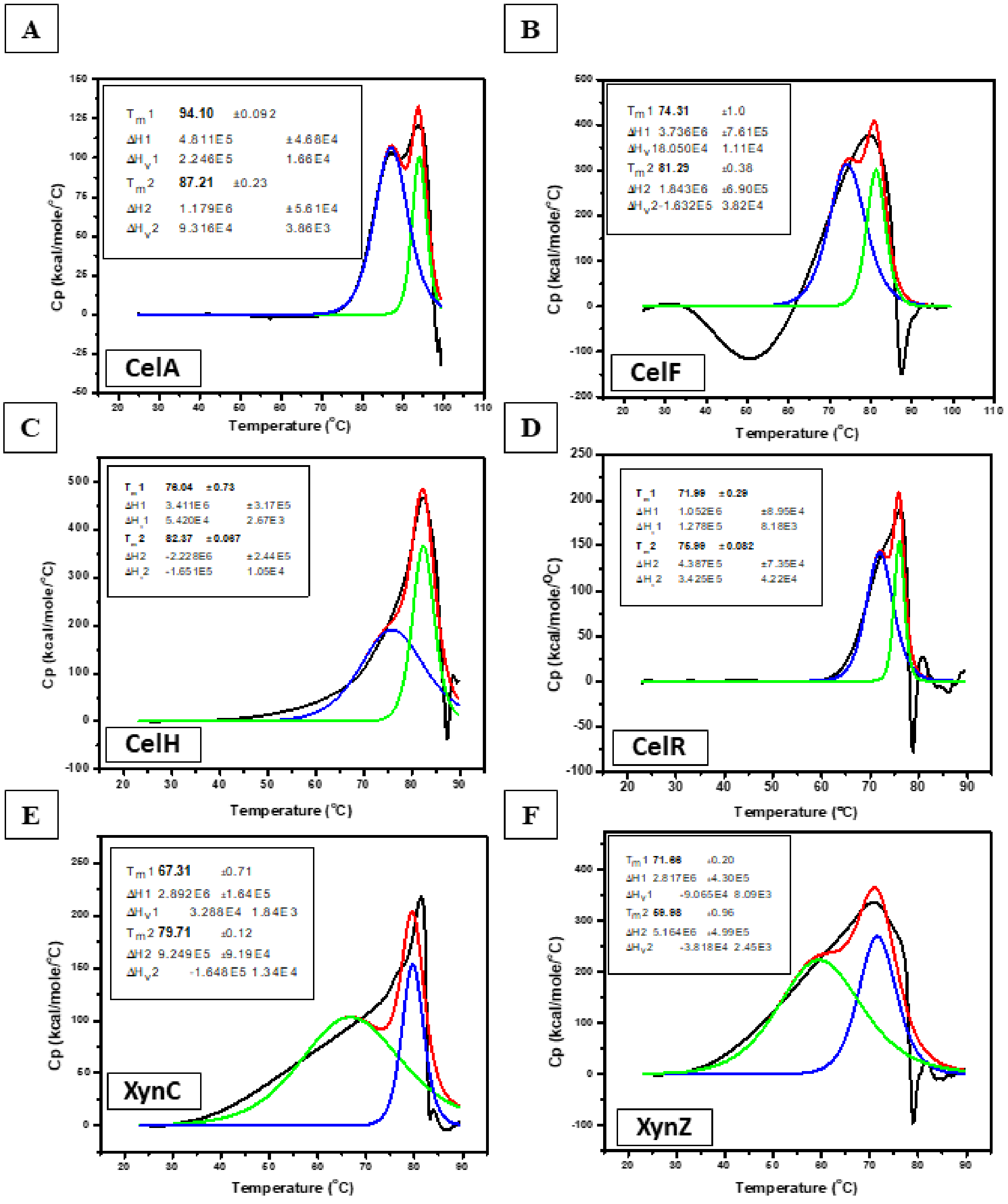


**Supplementary Figure S12**: Differential scanning calorimetry (DSC) data for the heating of six of the seven enzyme-bearing Doc domains selected for this study (data for the seventh is shown in the main manuscript, Figure 2F). The DSC data helps to establish the highly thermostable natures of the enzymes, as well as the presence of multiple autonomously unfolding domains.

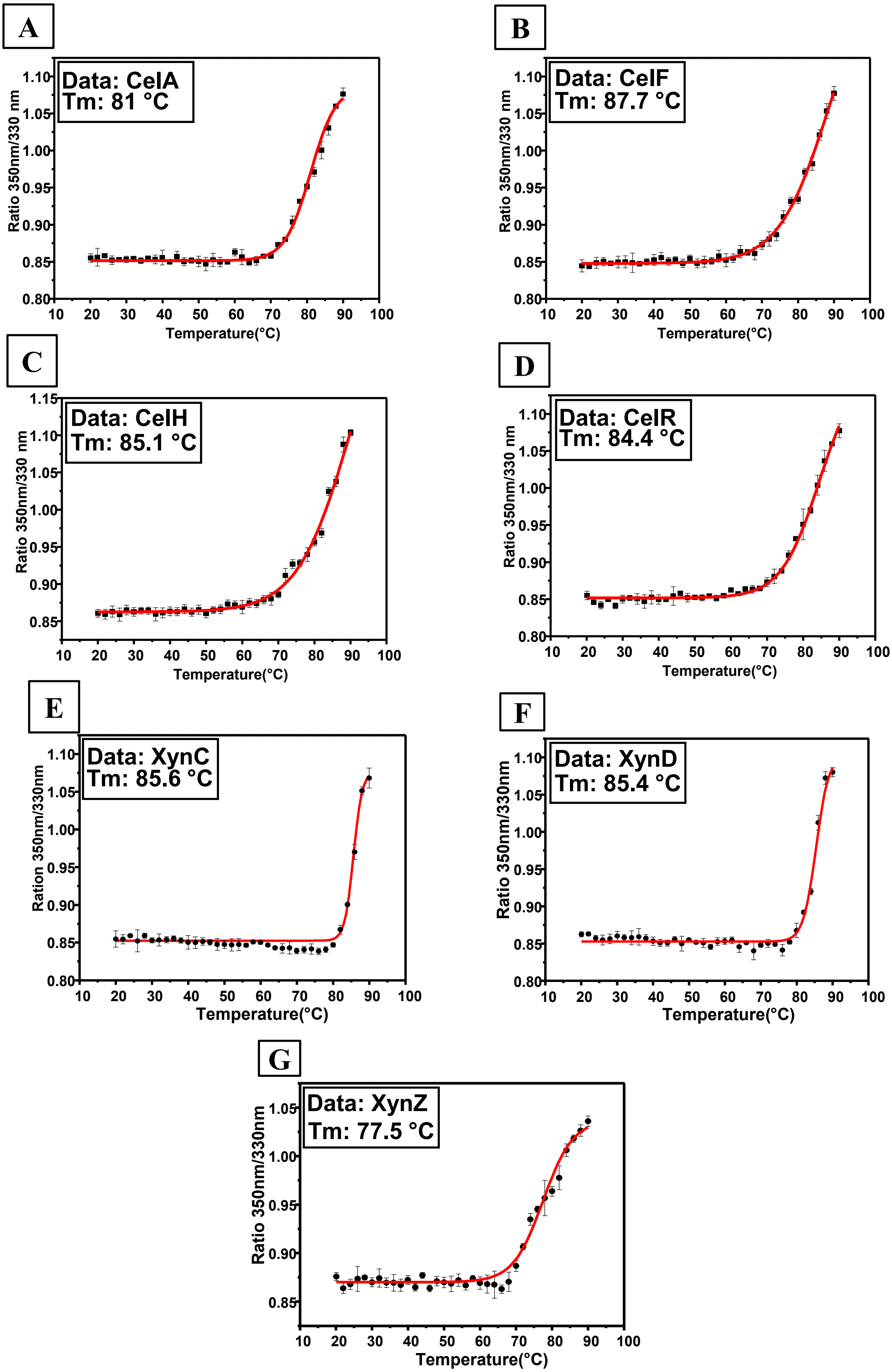


**Supplementary Figure S13**: Tryptophan fluorescence (350/330 nm ratios) based examination of the cooperative (thermal) unfolding of the selected enzyme-bearing Doc domains.

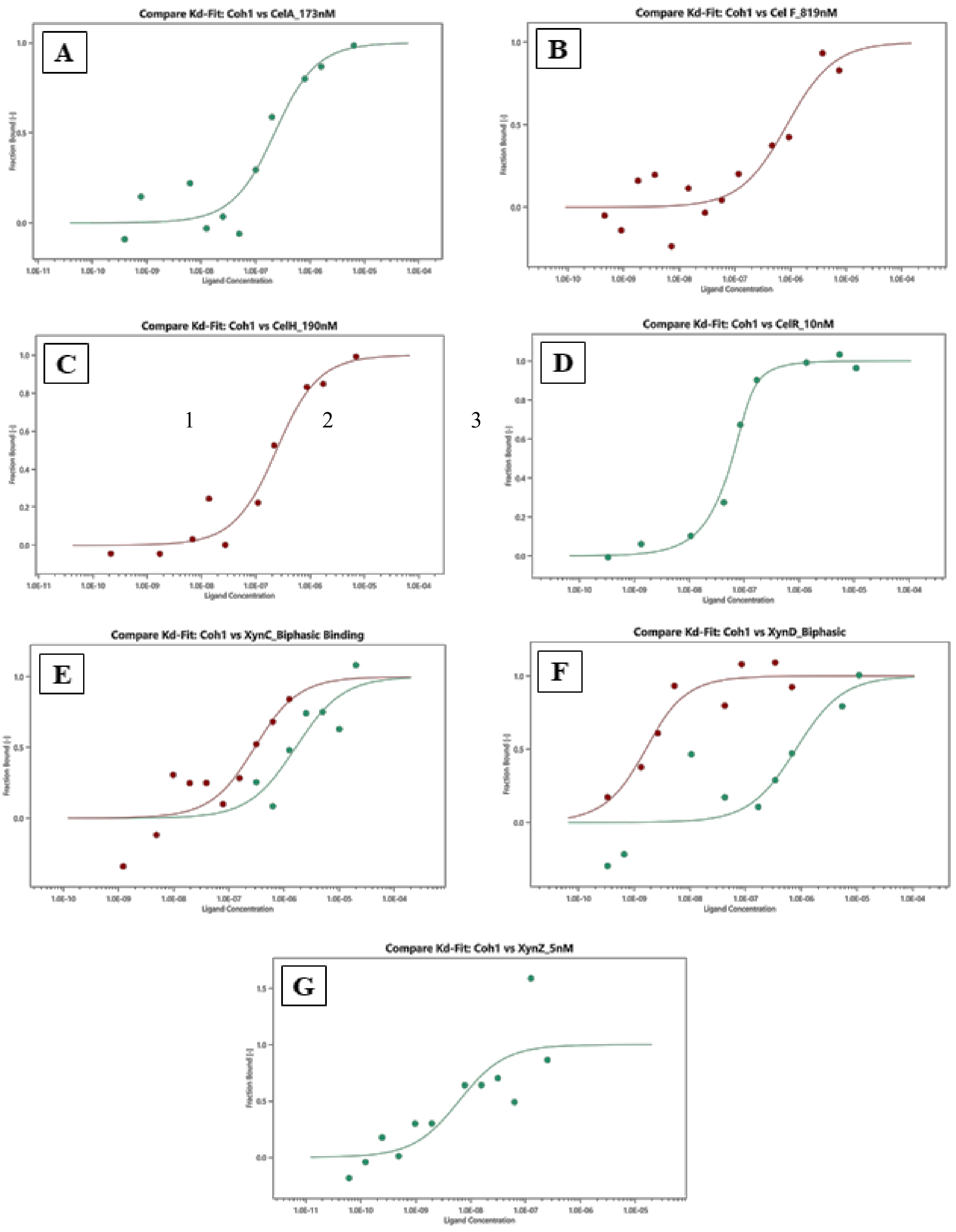


**Supplementary Figure S14**: Quantitative interaction analysis of Coh1 with selected enzyme-bearing Doc domains, based on data from microscale thermophoresis (MST). (A) CelA (B) CelF (C) CelH (D) CelR (E) XynC (F) XynD and (G) XynZ. X axis represents ligand concentration in M whereas Y axis represents fraction bound as described in methods; this is mentioned here because we are unable to improve the figure quality.

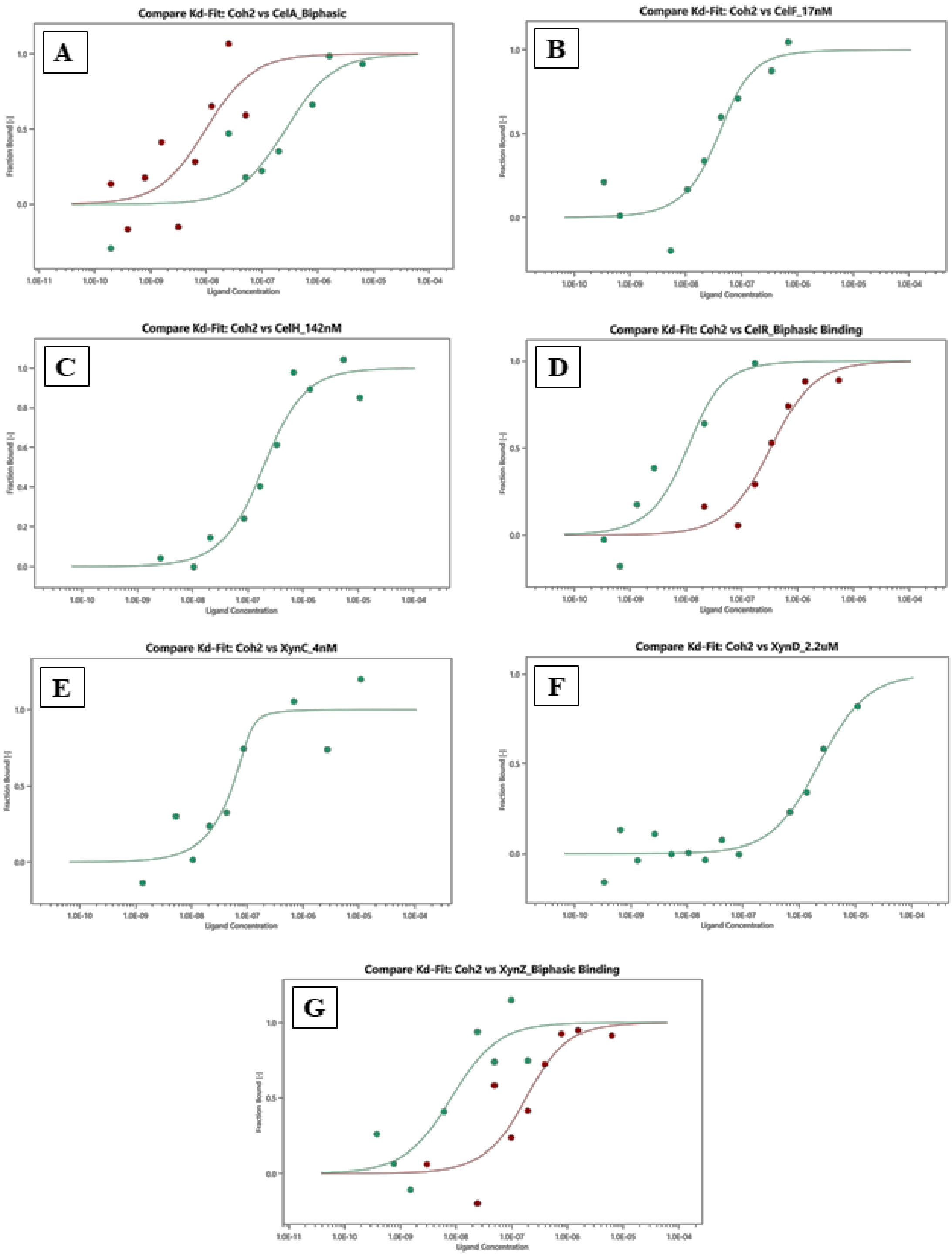


**Supplementary Figure S15**: Quantitative interaction analysis of Coh2 with selected enzyme-bearing Doc domains, based on data from microscale thermophoresis (MST). (A) CelA (B) CelF (C) CelH (D) CelR (E) XynC (F) XynD and (G) XynZ. X axis represents ligand concentration in M whereas Y axis represents fraction bound as described in methods; this is mentioned here because we are unable to improve the figure quality.

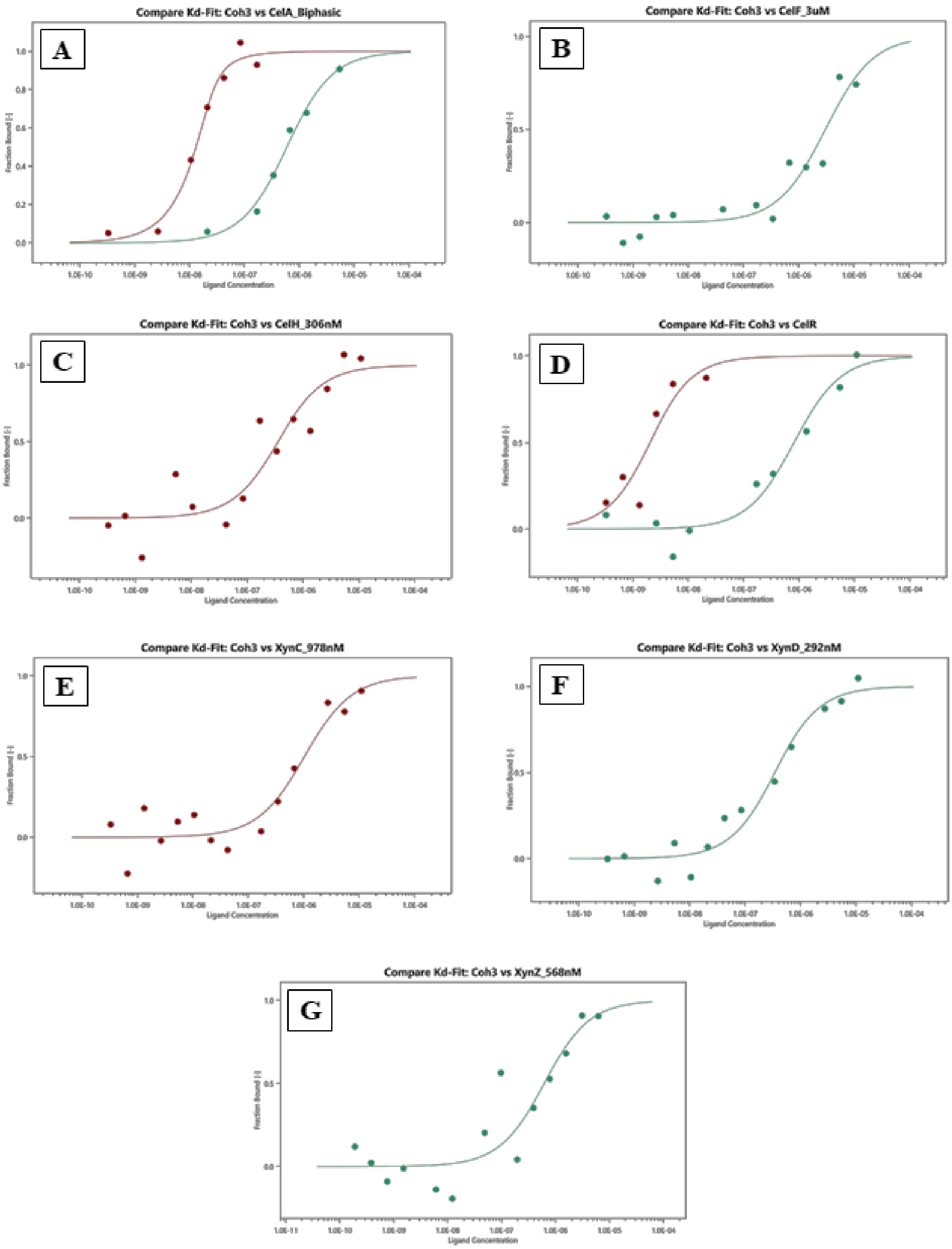


**Supplementary Figure S16**: Quantitative interaction analysis of Coh3 with selected enzyme-bearing Doc domains, based on data from microscale thermophoresis (MST). (A) CelA (B) CelF (C) CelH (D) CelR (E) XynC (F) XynD and (G) XynZ X axis represents ligand concentration in M whereas Y axis represents fraction bound as described in methods; this is mentioned here because we are unable to improve the figure quality.

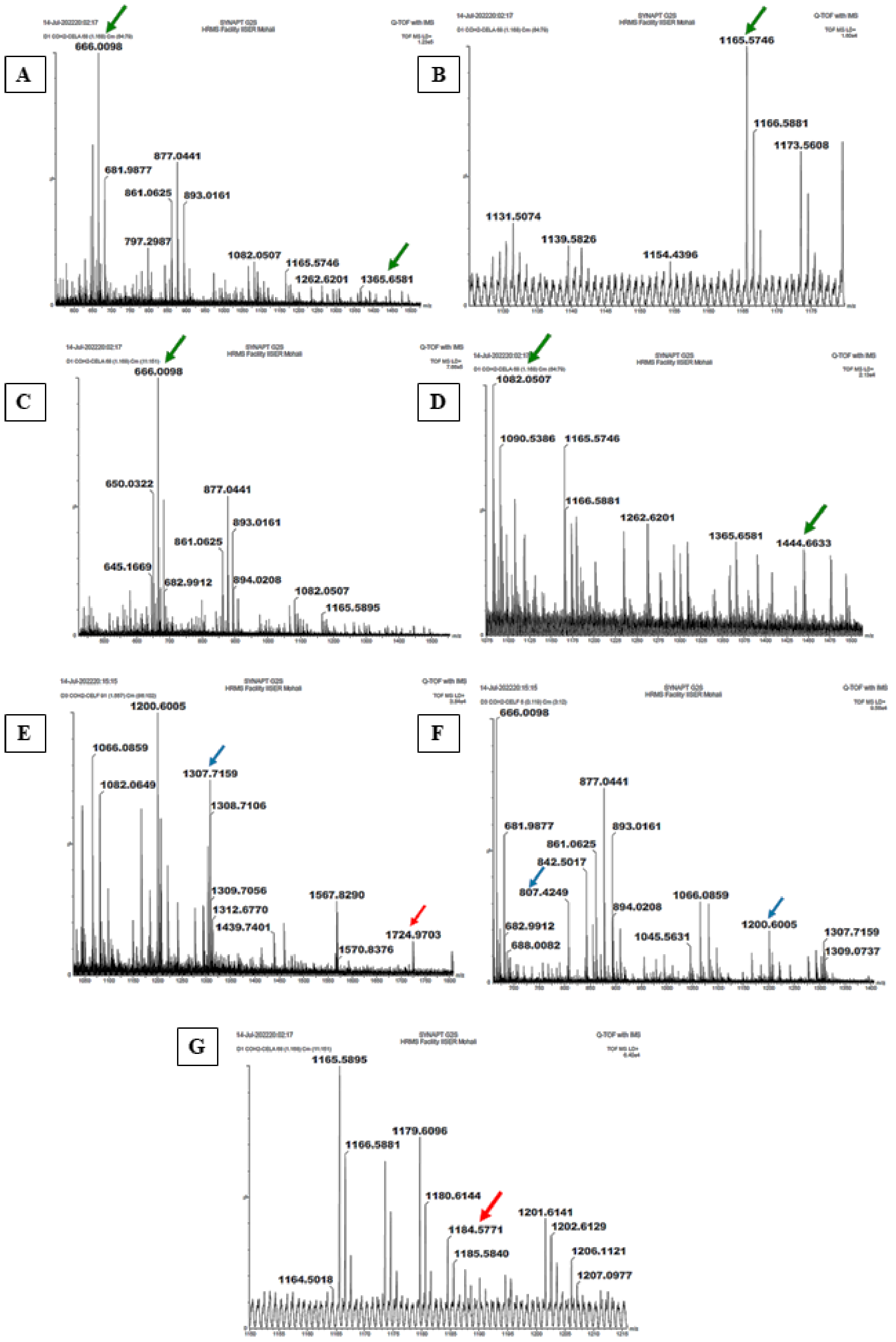


**Supplementary Figure S17**: Mass spectrometric confirmation of the identities of the complexes of interacting Coh and enzyme-bearing Doc domains observed in native PAGE, through peptide mass fingerprinting. Blue, green and red arrows indicate observations of peptides derived from CelF, CelA, and Coh2 respectively. Panels (A) to (D) show mass spectral data for Coh2 and CelA interaction, whereas, panels (E) to (G) show mass spectral data for Coh2 and CelF interaction.

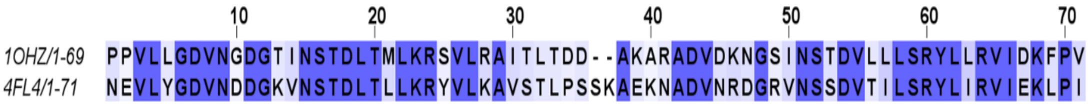


**Supplementary Figure S18**: Pairwise sequence alignment of Doc domains from the PDB structures, 1OHZ and 4FL4, made using UniProt align (Clustal Omega) and showing the high level of conservation of sequence. It may be noted that the Doc domain sequences shown in the above figure are from determined crystal structures, and not from any of the seven enzymes studied.

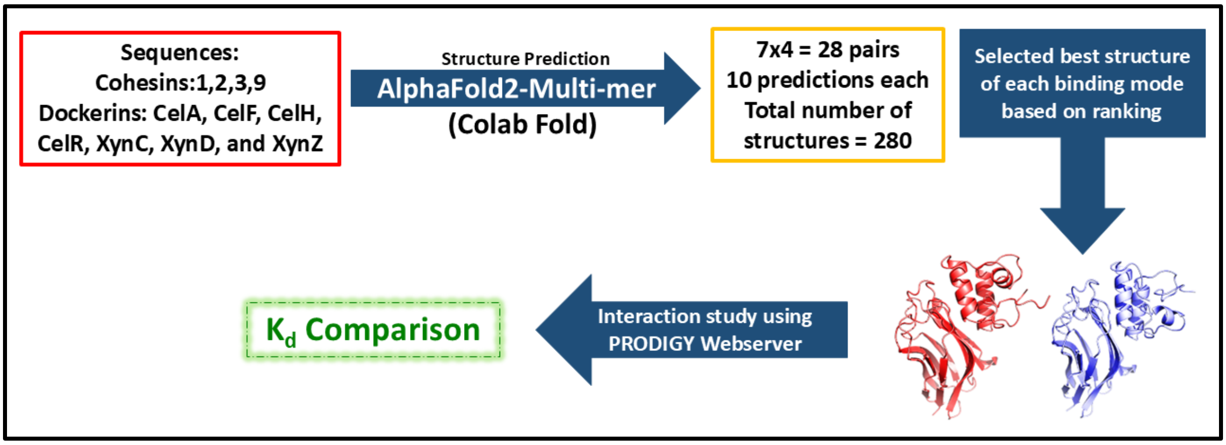


**Supplementary Figure S19**: Workflow diagram for structure prediction and interaction analyses using bioinformatic tools. Ranking of the model was based on the output from Colab Fold (based on pLDDT, pTM and ipTM scores).
